# A Chromatin Accessibility Atlas of the Developing Human Telencephalon

**DOI:** 10.1101/811620

**Authors:** Eirene Markenscoff-Papadimitriou, Sean Whalen, Pawel Przytycki, Reuben Thomas, Fadya Binyameen, Tomasz J. Nowakowski, Stephan J. Sanders, Matthew W. State, Katherine S. Pollard, John L. Rubenstein

## Abstract

Gene expression differs between cell types and regions within complex tissues such as the developing brain. To discover regulatory elements underlying this specificity, we generated genome-wide maps of chromatin accessibility in eleven anatomically-defined regions of the developing human telencephalon, including upper and deep layers of the prefrontal cortex. We predicted a subset of open chromatin regions (18%) that are most likely to be active enhancers, many of which are dynamic with 26% differing between early and late mid-gestation and 28% present in only one brain region. These region-specific predicted regulatory elements (pREs) are enriched proximal to genes with expression differences across regions and developmental stages and harbor distinct sequence motifs that suggest potential upstream regulators of regional and temporal transcription. We leverage this atlas to identify regulators of genes associated with autism spectrum disorder (ASD) including an enhancer of *BCL11A*, validated in mouse, and two functional *de novo* mutations in individuals with ASD in an enhancer of *SLC6A1*, validated in neuroblastoma cells. These applications demonstrate the utility of this atlas for decoding neurodevelopmental gene regulation in health and disease.

**Summary:** To discover regulatory elements driving the specificity of gene expression in different cell types and regions of the developing human brain, we generated an atlas of open chromatin from eleven dissected regions of the mid-gestation human telencephalon, including upper and deep layers of the prefrontal cortex. We identified a subset of open chromatin regions (OCRs), termed predicted regulatory elements (pREs), that are likely to function as developmental brain enhancers. pREs showed regional differences in chromatin accessibility, including many specific to one brain region, and were correlated with gene expression differences across the same regions and gestational ages. pREs allowed us to map neurodevelopmental disorder risk genes to developing telencephalic regions, and we identified three functional *de novo* noncoding variants in pREs that alter enhancer function. In addition, transgenic experiments in mouse validated enhancer activity for a pRE proximal to *BCL11A*, showing how this atlas serves as a resource for decoding neurodevelopmental gene regulation in health and disease.

## Introduction

The development of the human telencephalon, the seat of cognition and consciousness, requires the stepwise generation of regions and cell types, long-distance migrations of cells and axons, and the formation of precise connections (J. Rubenstein and Rakic 2013). This spatiotemporal precision is mirrored in gene expression patterns across the telencephalon, which are orchestrated by transcription factors (TFs) binding to diverse classes of proximal and distal regulatory elements. Genomic regulatory elements (REs) play an important role in forebrain development (Visel et al. 2007, 2013; Dickel et al. 2018), and identifying REs specific to cell types and brain regions is an important step towards elucidating the transcriptional mechanisms underlying human brain development and interpreting genetic risk variants associated with neurodevelopmental disorders.

Experiments in mice have illustrated the remarkable specificity and importance of distal enhancers that are evolutionarily conserved in humans during mammalian telencephalon development (Silberberg et al. 2016; Pattabiraman et al. 2014; Visel et al. 2007, 2013; Erwin et al. 2014; Dickel et al. 2018). Advancements in single-cell techniques have recently allowed unprecedented analysis of cell types in the developing human brain (Nowakowski et al. 2017), and innovative computational approaches have been developed to predict gene expression changes that drive cell type specification (La Manno et al. 2018). Longitudinal transcriptomic analyses of the human brain have shown that early to mid-fetal development is a critical time for diversification, exhibiting high regional variation in gene expression which declines in late fetal development as well as postnatally (M. Li et al. 2018).

Chromatin accessibility is a common feature of REs (de la Torre-Ubieta et al. 2018). Mapping the dynamics of the chromatin accessibility landscape that accompanies, and likely determines, gene expression changes during development is possible with techniques like ATAC-seq (Buenrostro et al. 2015). A recent study used this technique to map chromatin accessibility changes across human cortical neuron differentiation and identify neural progenitor enhancers (de la Torre-Ubieta et al. 2018). Thus, ATAC-seq maps can be improved by annotating a subset of open chromatin regions (OCRs) most likely to regulate gene expression. Our work expands on this study by 1) assaying anatomically distinct cortical regions, including sub-cortical regions of the telencephalon that have not been analyzed previously, 2) tracking changes between early and late mid-fetal development, 3) dissecting differences in accessibility between cortical laminae, and 4) identifying thousands of candidate enhancers specific to these processes.

The resulting map of chromatin accessibility in the developing human brain adds a greater degree of spatial and temporal resolution to previous maps (de la Torre-Ubieta et al. 2018; Roadmap Epigenomics Consortium et al. 2015). This higher resolution is important for several reasons. First, many developmental genes are broadly expressed in one or more tissues and stages, but each component of their expression pattern may be regulated by distinct enhancers. Intersecting genetic risk alleles with a spatio-temporal open chromatin atlas could implicate specific brain regions or developmental periods in pathological processes. Such an approach has already been successful in the immune system (Farh et al. 2015). Furthermore, if disease risk is restricted to a subset of a gene’s expression domains, testing enhancers unique to that subset for their disease association will increase statistical power relative to testing broader sets of enhancers. We expect this increase in resolution and power will be critical for identifying causal variants in loci that have been linked to disease using common variants, as well as distinguishing rare and *de novo* variants in cases versus controls.

We generated an atlas of the chromatin accessibility landscape of the mid-fetal human telencephalon. ATAC-seq was performed on nuclei extracted from samples of six cortical regions and the basal ganglia anlage (the three ganglionic eminences) dissected from intact specimens. We identified statistically significant OCRs, generated a list of predicted regulatory elements (pREs) from these OCRs, and validated enhancer activities using luciferase transcription and transgenic enhancer assays. Spatiotemporal differences in gene expression across the developing telencephalon were associated with differential chromatin accessibility at pREs. We further identified pREs that may drive laminar gene expression differences and the development of upper and deep layers of the cortex. Motif analysis uncovered TFs that may bind pREs and drive enhancer specificity. Finally, we integrated *de novo* variants identified from whole genome sequencing (WGS) of autism spectrum disorder (ASD) cases and identified functional mutations in a pRE located in an intron of ASD risk gene *SLC6A1*. Thus, in addition to enumerating open chromatin regions and candidate enhancers that drive regional and cellular identity in the developing human telencephalon, this atlas also provides a framework for predicting the cell types and brain regions impacted by noncoding genetic risk variants during development.

## Results

### Identifying open chromatin regions in the mid-gestation human telencephalon

We performed ATAC-seq on five fresh, intact samples of mid-gestation (14-19 gestation weeks [gw], measured from the last menstrual period) human fetal telencephalon, including six regions dissected from cortical anlage: dorsal lateral prefrontal cortex (PFC), motor, primary somatosensory cortex (S1), temporal cortex, parietal cortex, and primary visual cortex (V1); and three subcortical regions (basal ganglia anlage): medial, lateral, and caudal ganglionic eminences (MGE, LGE, CGE) (Figure 1A). Based on mouse fate mapping experiments, the MGE generates pallidal neurons and interneurons that integrate into the cortex and striatum, the LGE generates striatal medium spiny neurons, and the CGE generates subpallial amygdala neurons and cortical interneurons (J. L. R. Rubenstein and Campbell 2013). The regional dissections spanned the full thickness of the telencephalic wall, from the ventricular zone to the meninges. Thus, all samples included progenitors and immature neurons, glia, blood vessels, and meninges. Twenty-five ATAC-seq libraries were generated as described (Buenrostro et al. 2015) and sequenced using Illumina Hiseq 2500 to an average depth of 100 million reads per sample. Reads were mapped to the human genome build GRCh37 using the ENCODE pipeline (Lee et al. 2016) and were highly correlated with published DNase hypersensitivity loci (Roadmap Epigenomics Consortium et al. 2015) (mean 58.8% within universal DHS peaks; Table S1A).

**Figure 1:**
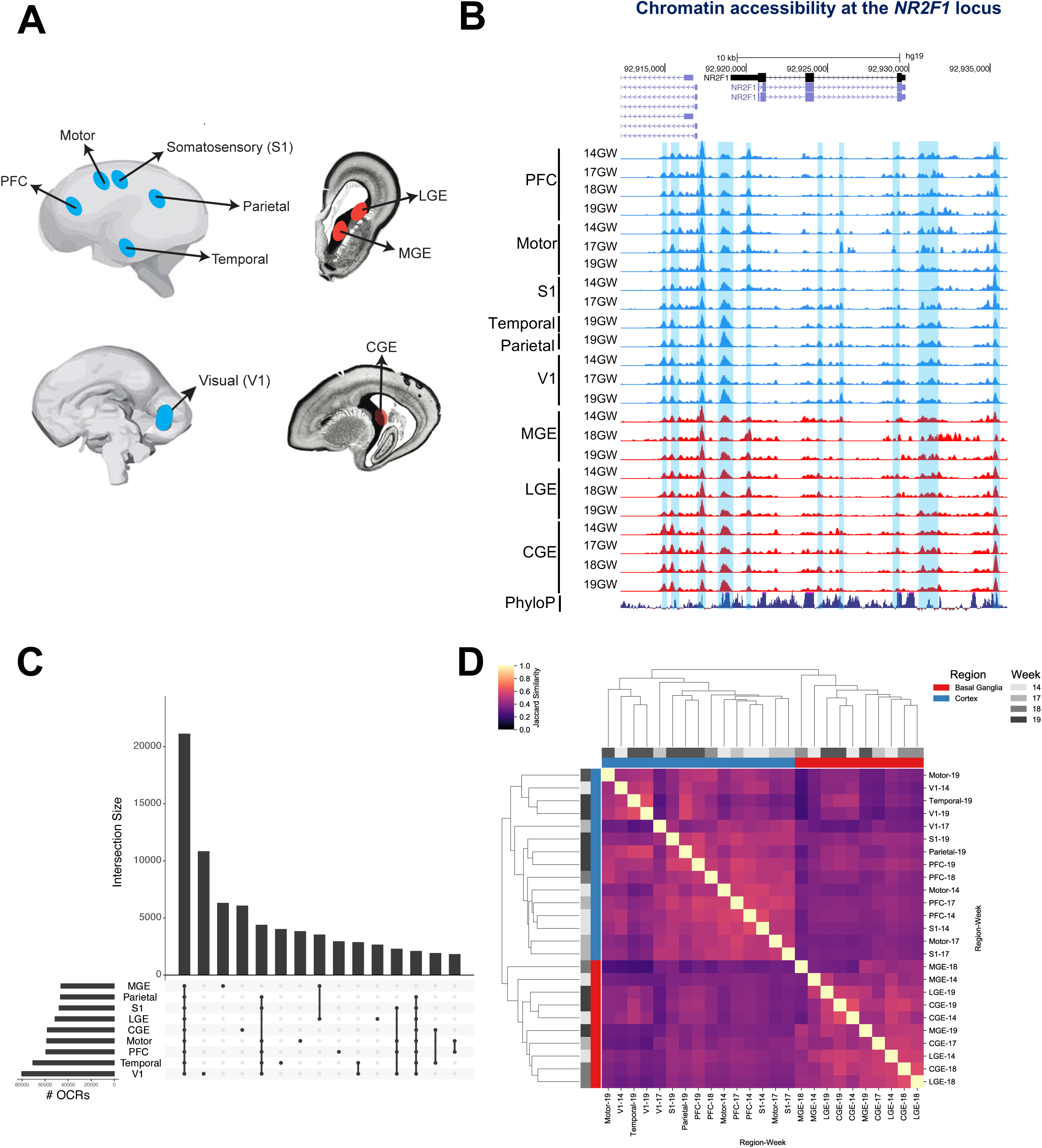
Defining open chromatin regions (OCRs) in the fetal human telencephalon. 1A) Schematic of fetal tissue collection. 1B) ATAC-seq reads from all fetal samples collected mapped to the *NR2F1* locus. Blue shaded regions are OCRs. Y axis scale is 0 to 10. 1C) OCR intersections and # OCRs pooled across samples for each region. 1D) Heat map of jaccard similarity of OCRs in all samples.

The ENCODE pipeline identified statistically significant open chromatin regions (OCRs) in twenty-four ATAC-seq libraries that passed stringent quality filtering (Figure 1B, Methods). We obtained a median of 36,325 high-confidence OCRs per sample (Table S1A), totaling 130,131 unique OCRs merged across all telencephalic regions. Seventeen percent of OCRs are shared between all assayed regions, while 3.7% are shared between all assayed cortical regions. This leaves many OCRs that are specific to one region; primary visual cortex, MGE, and CGE in particular (Figure 1C). Clustering samples by OCR similarity shows two main groups representing the cortex and basal ganglia (Figure 1D). Within cortex and basal ganglia, samples cluster by age rather than subregion.

### Predicting developmental brain enhancers

Chromatin accessibility is elevated at known forebrain enhancers and reduced at enhancers for other tissues: examples of a forebrain enhancer (hs433) and a limb enhancer (hs72) are shown in Figures 2A and 2B. To identify which OCRs are likely to function as neurodevelopmental enhancers, we developed a model using a machine learning algorithm trained on candidate enhancers tested in vivo using transient transgenic mouse reporter assays, as well as a diverse set of features (Methods). Specifically, we included all human sequences from the VISTA Enhancer Browser database (Visel et al. 2007) with strong embryonic brain enhancer activity as true positives, compared to sequences that showed activity in non-brain tissues or weak/no activity in VISTA as true negatives. Each positive and negative sequence was annotated with functional genomics data and the sub-sequences (k-mers) it contains. The algorithm then learned from these features how to distinguish the positive brain enhancer regions from the negative control regions (Figure S2).

**Figure 2:**
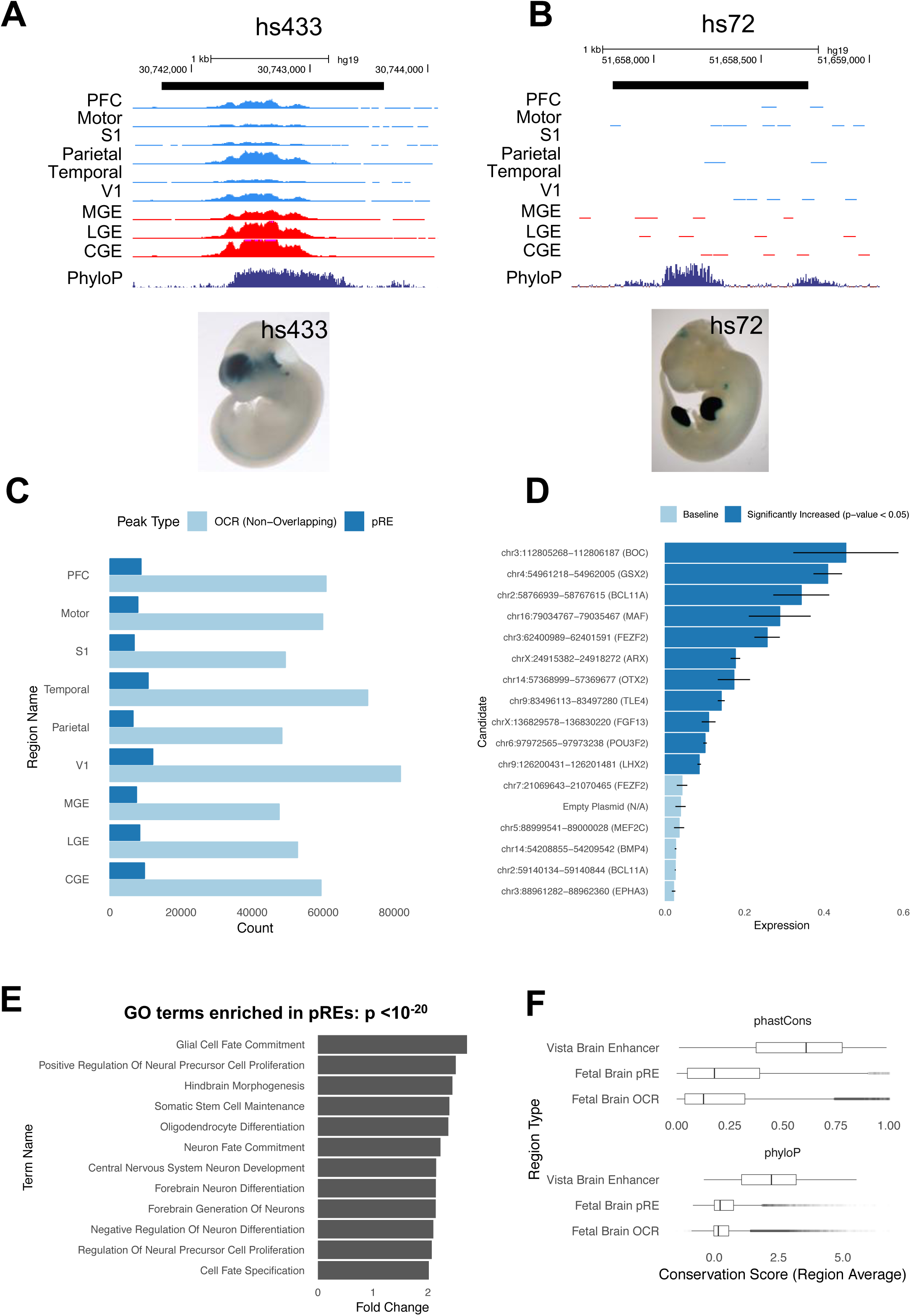
Predicting neurodevelopmental enhancers. 2A) ATAC-seq reads, combined across multiple samples per brain region, at VISTA brain enhancer hs433. Y axis scale is 0 to 500. E11.5 enhancer transgenic mouse hs433 is shown and lacZ is stained in blue (from Vista enhancer website). 2B) ATAC-seq reads, combined across multiple samples per brain region, at limb VISTA enhancer 72. Y axis scale is 0 to 500. E11.5 enhancer transgenic mouse hs72 is shown and lacZ expression is stained in blue (from Vista enhancer website). 2C) Fraction of OCRs that are pREs in each brain region. 2D) Mean firefly luciferase levels in human neuroblastoma cells, normalized to Renilla. Fifteen pREs (chromosomal locations and nearest gene indicated) were cloned upstream of a minimal promoter and firefly luciferase ORF in the pGL4.23 vector. Dark blue bars indicate pREs with increased luciferase signal activity compared to empty vector, error bars indicate standard error across four replicate experiments. *ARX* VISTA enhancer hs119 is included as a positive control for comparison. 2E) Gene ontology terms enriched in pREs with GREAT analysis, complete list in Table S2C. 2F) Phastcons and PhyloP scores (100-way alignments, vertebrate conservation) of pREs, OCRs, and VISTA enhancers.

Applying the resulting model to all 130,131 OCRs, we identified a subset of 19,151 predicted regulatory elements (pREs) that do not overlap with coding regions or promoters (Table S2A) and comprise 18.4% of all OCRs (Figure 2C, median 6,918 pREs per brain region). We expect these pREs consist primarily of distal enhancers, though other *cis*-regulatory elements may be also be included. We tested the ability of 15 pREs with high prediction scores (>0.85) to regulate transcription using a transfection assay with a luciferase readout. Ten pREs (66%) were validated by this assay in a mitotically active human brain derived BE(2)-C cell line (Figure 2D).

Gene ontology analysis of the nearest genes to pREs using GREAT shows enrichment of neuro-developmental terms such as “central nervous system development”, “neuron differentiation”, and “forebrain generation of neurons” (Figure 2E), while no terms are enriched for genes proximal to non-pRE OCRs. pREs are more highly conserved than OCRs, but not as conserved as the training set of VISTA enhancers, which were originally selected for having exceptionally high conservation scores (Visel et al. 2007) (Figure 2F).

### Identifying region-specific predicted enhancers

We next associated differences in gene expression with regional differences in chromatin accessibility at pREs. First, we used published single-cell RNA-seq data from 14-19gw MGE, PFC, and V1 samples (Nowakowski et al. 2017). A subset of the specimens used to generate the RNA-seq data were also used to generate our ATAC-seq libraries and thus are well-matched. We then identified 1,800 differentially expressed (DE) genes pairwise between the MGE, PFC, and V1. pREs with differential chromatin accessibility were highly enriched for DE genes (odds ratios > 4, Figure 3A). We identified 510 differentially accessible pREs that were associated with DE genes between cortex (PFC or V1) and MGE (Table S3A,B). For example, MBIP is more highly expressed in MGE compared to PFC and V1 (3-fold change, q-value 1.2*10^-16^), while *KCNQ3* has the opposite expression pattern (4-fold change, q-value 1.3*10^-13^). Both *MBIP* and *KCNQ3* have accompanying differences in chromatin accessibility at proximal pREs (Figures 3B,C). These results are supported by *in situ* hybridization experiments (Tucker et al. 2008; Diez-Roux et al. 2011) (Figure S3A).

**Figure 3:**
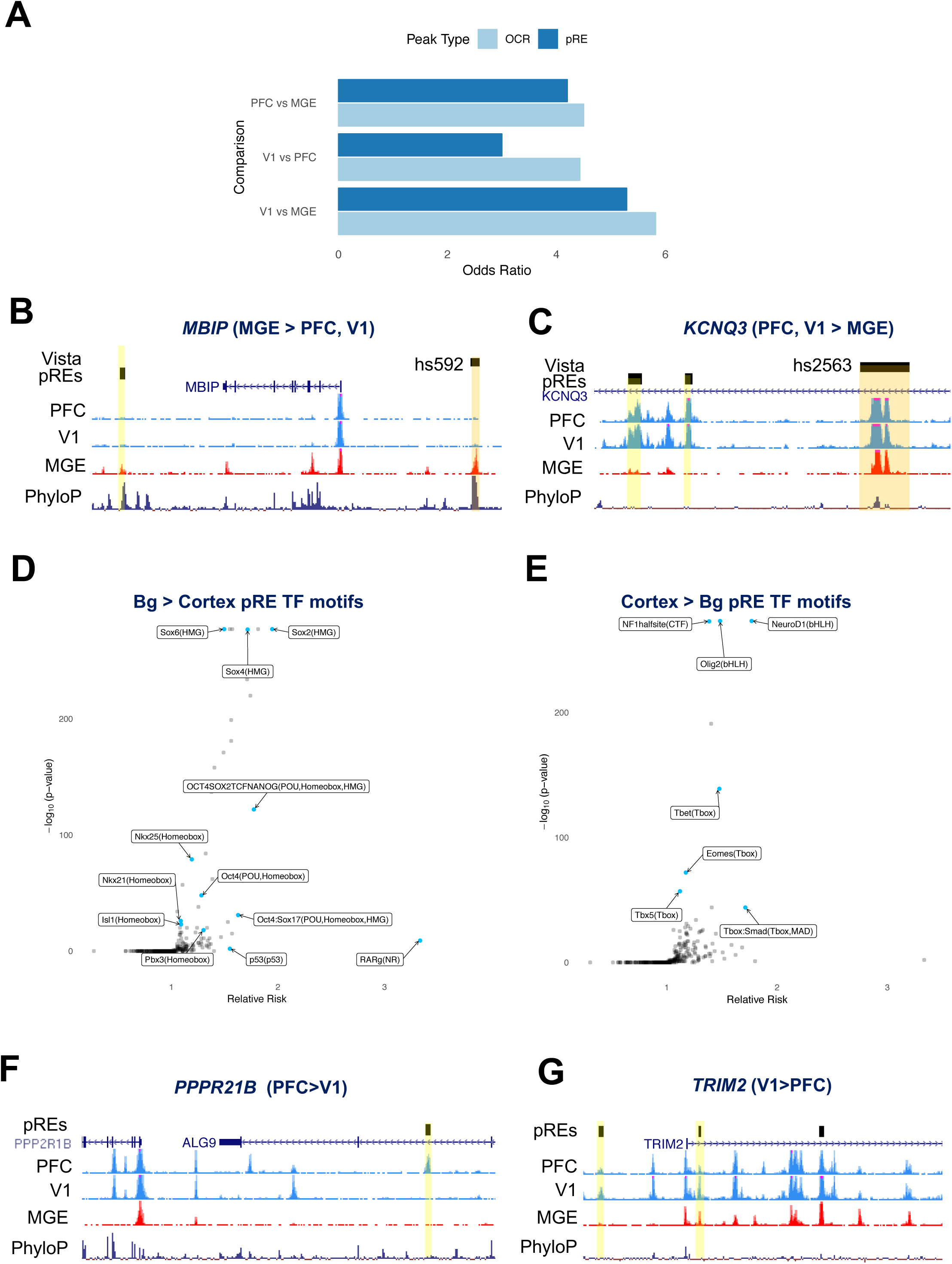
Regional differences in chromatin accessibility at pREs. 3A) Odds ratio for the co-occurence of genes that were differentially expressed and proximal to differentially accessible chromatin (100 kb), computed between pairs of brain regions and reported separately for OCRs and pREs. 3B,C) ATAC-seq reads combined from multiple samples in PFC, V1, and MGE. pREs are highlighted in yellow, VISTA enhancers in orange. The nearby genes are differentially expressed between the regions indicated. Y axis scale is 0 to 50 for ATAC-seq tracks. 3D) Effect size and significance of TF motifs enriched in basal ganglia-specific pREs compared to cortex-specific pREs. 3E) Effect size and significance of TF motifs enriched in cortex-specific pREs compared to basal ganglia-specific pREs. 3F,G) ATAC-seq reads combined from multiple samples in PFC, V1, and MGE. pREs are highlighted in yellow. The nearby genes are differentially expressed between the regions indicated. Y axis scale is 0 to 50 for ATAC-seq tracks.

Focusing on regional differences within the telencephalon, we identified 6,942 pREs (36.2%) that were cortex specific and 3,462 pREs (18%) that were basal ganglia specific (Table S2A). Here, we define “specific” as a pRE that has no statistically significant ATAC-seq signal in the other brain regions in our data. In order to explore potential upstream regulators of pREs, we looked for enrichment of TF binding motifs in these region-specific pREs. Basal ganglia specific pREs were enriched for several motifs including SOX TFs, with 9.9% containing the composite motif for OCT4-SOX2, a sequence that promotes MGE enhancer function in mice (Sandberg et al. 2018) (Figure 3D). Cortex specific pREs were enriched for motifs of cortical TFs that have well known functions in the developing cortex, including NEUROD1, OLIG2, NF1, and the T-BOX family members (e.g. TBR1 and TBR2/Eomes) (Figure 3E).

To discover elements likely driving rostral-caudal differences in gene expression within the cortex, we identified 79 pREs with differential accessibility between PFC and V1 that were proximal to genes differentially expressed in single-cell RNA-seq data between these same two regions (Table S3C). *PPP2R1B* is more highly expressed rostrally in PFC (1.2 fold change, q-value 0.019), while *TRIM2* is more highly expressed caudally in V1 (1.7 fold change, q-value 0.047) which is supported by *in situ* hybridization experiments (Diez-Roux et al. 2011) (Figure S3A). Both gene loci are proximal to differentially accessible pREs (Figures 3F and 3G).

To identify pREs that are accessible in only one brain region, we combined OCRs across timepoints within each region, identified OCRs exclusive to a single brain region, and filtered the OCR list to those that were also pREs (Figure S3B, Table S2A). The resulting 5,318 region specific pREs (27.7% of total) differed in their genomic distributions compared to other pREs, more often overlapping intronic and intergenic regions (Figure S3C).

### Identifying cortical developmental stage-specific pREs

Since dramatic differences are observed in gene expression over developmental time in the fetal brain (M. Li et al. 2018) and the ATAC-seq samples are seen to cluster by developmental age within a region (Figure 1D), we sought to quantify changes in chromatin accessibility between the early and late stages of mid-fetal cortical development. Combining samples from the PFC and motor cortex, we identified 1,330 pREs specific to 14gw and 3,559 specific to 19gw in frontal cortex samples. GREAT analysis to identify the nearest genes of the early mid-fetal specific pREs yielded gene ontology terms for processes occurring during early cortical development such as “cerebral cortex radially oriented cell migration” and “neural precursor cell proliferation”, while late mid-fetal specific pREs were enriched for processes occurring with neuronal maturation such as “axon extension”, “long term synaptic potentiation”, “neurotransmitter secretion”, and “neuron fate commitment” (Figures 4A, B, Table S4A).

**Figure 4:**
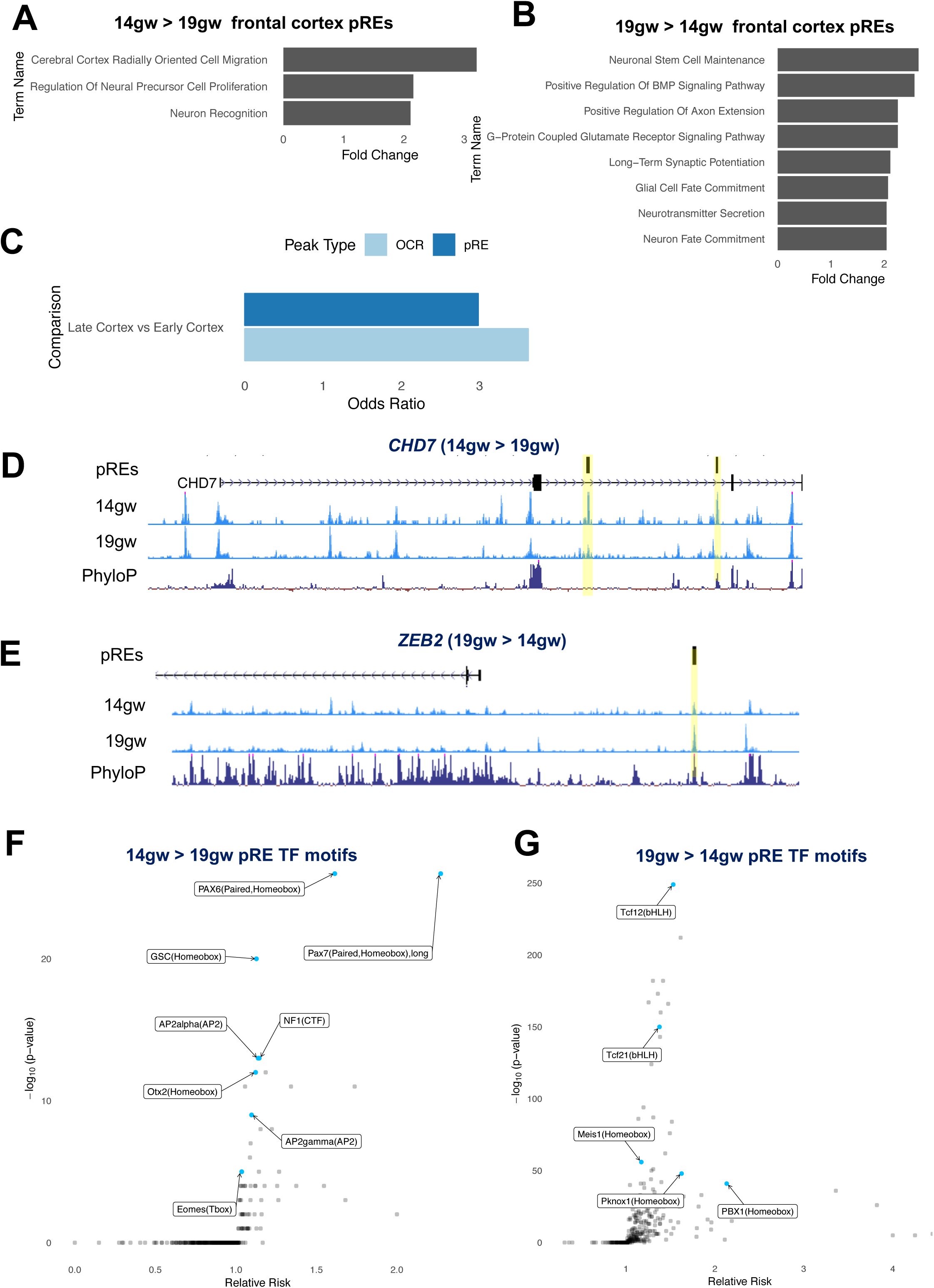
Temporal differences in chromatin accessibility at cortical pREs. 4A) GREAT analysis of functions associated with 14gw-specific pREs from frontal cortex tissues (combined PFC and motor samples). 4B) GREAT analysis of functions associated with 19gw-specific pREs from frontal cortex tissues. 4C) Genes differentially expressed in frontal cortex between 19gw and 14gw are enriched for 19gw- and 14gw-specific pREs and OCRs. 4D,E) ATAC-seq reads from 14gw and 19gw frontal cortex pooled samples. pREs are highlighted in yellow. The nearby genes are differentially expressed at 14gw and 19gw, respectively. Y axis scale is 0 to 50. 4F,G) Effect size and significance of TF motifs enriched in 14gw frontal cortex-specific pREs compared to 19gw-specific pREs, and vice versa.

To associate these differences in pREs with gene expression changes across mid-gestation, we integrated 14gw and 19gw pREs with single-cell RNA-seq data from age-matched samples (Nowakowski et al. 2017). 14gw- and 19gw-specific pREs in frontal cortex were enriched for genes differentially expressed between those ages (Figure 4C). We identified 203 age-specific pREs that were proximal to differentially expressed genes between 14gw and 19gw (Table S4B). For instance, the gene *CHD7* (Vissers et al. 2004), which is associated with CHARGE syndrome and is expressed in cortical progenitors in the ventricular zone (Figure S4A) (Diez-Roux et al. 2011), is more highly expressed in 14gw cortex compared to 19gw (2-fold change, q-value 0.0009). Our data shows that *CHD7* contains two intronic pREs that are specific to early but not late frontal cortex (Figure 4D). In contrast is *ZEB2*, which is expressed in immature cortical excitatory neurons (Seuntjens et al. 2009). *ZEB2* is more highly expressed in 19gw cortex (3-fold change, q-value 0.0027) and is likewise proximal to a pRE that is specific to late but not early frontal cortex development (Figure 4E).

Analysis of TF motifs in 14gw cortical pREs found enrichment of homeodomain TF motifs (e.g. PAX6), while motifs for BHLH TFs (e.g. TCF12) and homeobox TFs (PBX1) are enriched in 19gw cortical pREs (Figures 4F and 4G). Enrichment of these motifs may reflect the changing cellular makeup of the tissue: *PAX6* is specifically expressed in cortical progenitors in the ventricular zone which are more abundant early in cortical development, while *PBX1* is highly expressed in neurons that are more populous in later stages (O’Leary, Chou, and Sahara 2007; Golonzhka et al. 2015).

### Identifying putative enhancers for deep and superficial cortical projection neurons

Upper and deep layer excitatory projection neurons of the cortex have been predicted (Willsey et al. 2013; Parikshak et al. 2013) and shown (Velmeshev et al. 2019) to be altered in ASD. The regulatory elements that drive the specification of cortical projection neuron subtypes have not been identified in humans, although conserved enhancers have been studied in mice (Shim et al. 2012). We reasoned that candidate cortical neuron subtype enhancers could be identified by integrating our pREs with ATAC-seq data from upper and deep cortical layers.

We microdissected the upper and deep layers of the cortical plate of PFC from fresh 18gw and 19gw hemispheres along with whole PFC dissections that span from the ventricular zone to the pia (Figure 5A). We then performed ATAC-seq and sequenced libraries to an average depth of 55 million reads before processing. Three biological replicates for upper and deep layers of cortex passed quality control. Using these three samples, we identified 71,628 upper layer and 57,008 deep layer OCRs at an Irreproducible Discovery Rate of 10% (Figure 5B). Of these, 5,517 OCRs were specific to deep layers and 20,867 were specific to upper layers of PFC (Table S5A).

**Figure 5:**
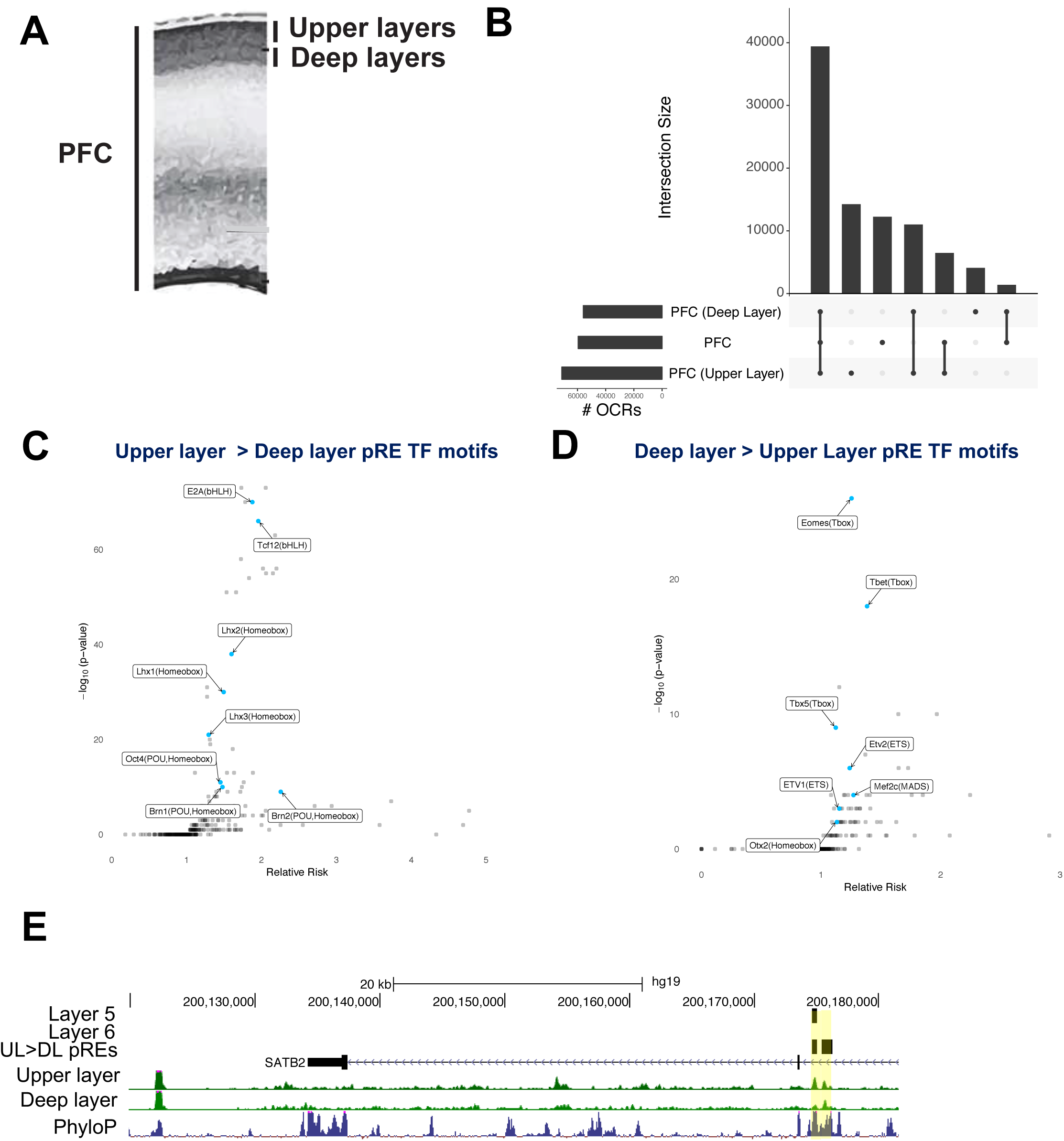
Laminar differences in chromatin accessibility in the developing PFC. 5A) Schematic of micro-dissection of upper and deep layers of cortex, for 18gw and 19gw ATAC-seq, and dissection of whole PFC, from germinal zone to cortical plate. 5B) OCR intersections and numbers of OCRs pooled across samples for upper layers, deep layers, and whole PFC. 5C) TF motifs enriched in upper layer-specific pREs. 5D) TF motifs enriched in deep layer-specific pREs. 5E) ATAC-seq reads combined from multiple samples from upper layer and deep layer microdissections of PFC at 18gw and 19gw, nearby upper layer gene *SATB2*. Highlighted in yellow are two pREs that are upper-layer specific (not OCR in deep layer). One of these two is also a conserved OCR in purified populations of layer 5 neurons in Rbp4-CRE transgenic mice, but not a conserved layer 6 neuron OCR in Ntsr1-CRE transgenic mice. Y-axis scale is 0 to 50.

To investigate whether OCR differences between deep and superficial PFC layers reflect gene regulatory differences, we analyzed TF motifs in the layer-specific OCRs that overlapped our original pREs. TF motif analysis of 2,382 upper layer-specific pREs showed enrichment for LHX (LIM HOMEODOMAIN), BRN (POU HOMEODOMAIN), and BHLH family motifs (Figure 5C).

These motifs are bound by transcription factors whose expression is elevated in superficial layers of the mouse and human cortex, including LHX2, LHX5, BRN1, BRN2, and BHLHB5 (Sestan and State 2018). Similarly, motif analysis of 461 deep layer-specific pREs showed enrichment for T-box and ETS family TFs (Figure 5D). These motifs are bound by TBR1, ETV1 and ETV2, transcription factors whose expression is elevated in deep layers (Sunkin et al. 2013).

To identify pREs that may be driving laminar differences in gene expression, we linked pREs with markers of upper and deep layer neurons clustered by t-SNE analysis of single-cell RNA-seq of human midfetal cortex (Nowakowski et al. 2017). Upper and deep layer neuron marker genes were enriched for proximity to upper and deep layer-specific pREs (odds ratios 6.44 and 2.22 respectively, p-value < 0.05). For example, the superficial layer marker gene *ADCYAP1R1* is proximal to an upper layer-specific pRE, while deep layer marker gene *PPFIA2* has a proximal pREs with deep layer-specific chromatin accessibility (Figure S5A,B).

At 18gw, the upper cortex layers consist of layer 2-5 neurons while deep layers consist of layer 5-6 and subplate neurons (Nowakowski et al. 2016). To assign pREs to distinct cortical neuron subtypes, we used cells purified from mouse CRE lines that label layer 5 subcortical projection neurons and layer 6 corticothalamic projection neurons using Rbp4-CRE and Ntsr1-CRE BAC transgenic mice, respectively (Gong et al. 2003). We crossed the CRE mice to Intact GFP reporter mice (Mo et al. 2015) and performed ATAC-seq on FACS-purified nuclei from prefrontal cortex at postnatal day 2 (Methods). After calling ATAC-seq peaks and lifting over to the human genome, we found 7,124 (12.5%) and 7,025 (12.3%) of human midfetal deep layer OCRs intersect mouse layer 5 and layer 6 OCRs, respectively. Similarly, 393 (7.5%) and 413 (7.9%) of human mid-fetal deep layer pREs overlap mouse layer 5 and layer 6 OCRs, respectively (Table S5A). Some conserved pREs are differentially accessible across layers 5 and 6. For example, we identified an upper layer pRE for the canonical upper layer gene *SATB2* that overlaps layer 5 but not layer 6 neuron OCRs identified in neonatal mouse (Figure 5E).

### Candidate regulatory elements for neurodevelopmental disorder genes

*BCL11A* is a gene associated with developmental delay and ASD (Sanders et al. 2015) that encodes a transcriptional regulator in brain and blood cells. There are 23 pREs within one megabase of the *BCL11A* locus with diverse patterns of accessibility across telencephalic regions (Figure 6A). We validated one of these for enhancer function *in vivo*, choosing a pRE based on high (>0.85) enhancer prediction score and luciferase transcription enhancer activity (Figure 2D, labeled *BCL11A*). This pRE is an OCR in the basal ganglia as well as cortical regions (Figure 6A). We cloned the pRE into the CT2IG GFP reporter vector (Silberberg et al. 2016) and generated transgenic mice by pronuclear injection. Like the endogenous *BCL11A* gene, two transgenic founders showed GFP expression in cortex, striatum, and hippocampus at postnatal day 2 (Figure 6B, Figure S6A).

**Figure 6:**
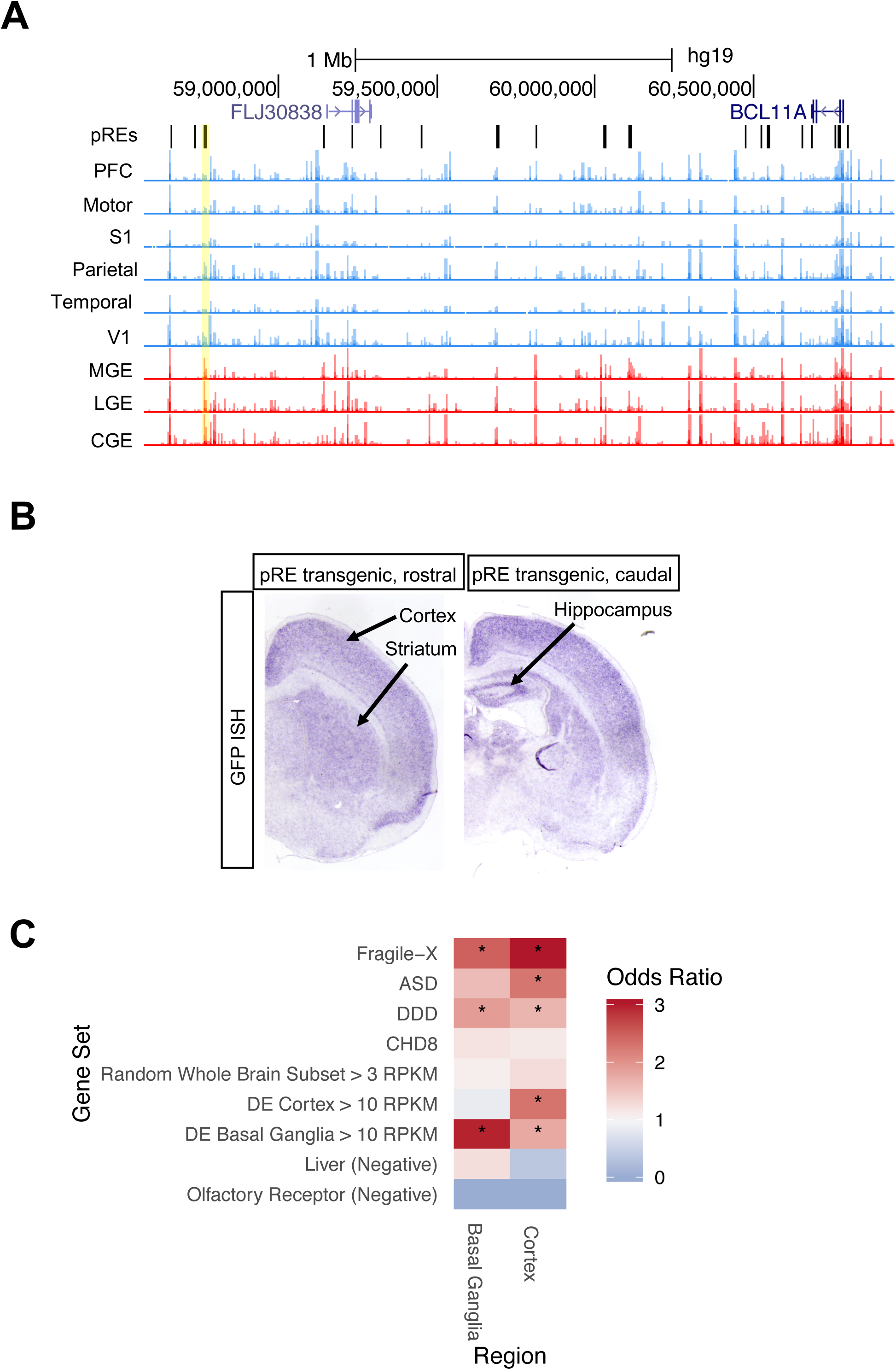
Identifying pREs that regulate neurodevelopmental disorder genes. 6A) ATAC-seq reads, combined across multiple samples per brain region, at pREs proximal to *BCL11A* on chromosome 2. The yellow shaded pRE was positive in luciferase assay (FIgure 2D) and was tested by transgenic enhancer mouse assay. Y axis scale is 0 to 500. 6B) Postnatal day 2 coronal sections of *BCL11A* enhancer transgenic mouse with GFP reporter expression in purple (RNA *in situ* hybridization for GFP). Cortical, striatal, and hippocampal expression is indicated. 6C) Enrichment analysis of genes, modeling their presence or absence in various gene sets as the inverse of their most proximal TSS to region-specific OCRs. Gene sets included ASD genes (Sanders et al. 2015), biological targets of Fragile X Mental Retardation protein (FMRP) (Darnell et al. 2011), biological targets of ASD gene CHD8 (Cotney et al. 2015), and developmental delay disorder (DD) genes (Deciphering Developmental Disorders Study 2015). Negative controls included liver (Subramanian et al. 2005) and olfactory receptor (Rouillard et al. 2016) genes, as well as a size-matched subset of whole-brain expressed genes (RPKM > 3). To increase power for testing specificity, peaks were first aggregated into basal ganglia or cortex regions. Highly expressed genes (> 10 RPKM) that were differentially expressed in cortex were enriched for proximity to cortex RS OCRs but not basal ganglia. Similarly, highly expressed genes that were differentially expressed in basal ganglia were enriched for proximity to cortex and basal ganglia RS OCRs, though with a much higher odds ratio for basal ganglia.

Using a gene set enrichment analysis, we asked whether other neurodevelopmental disorder genes such as *BCL11A* are enriched for candidate REs that are region-specific. In contrast to our previous analyses using pREs (Figures 2-5), OCRs were used here due to the limited number of region-specific pREs. We started with sets of genes associated with neurodevelopmental disorders including ASD (Sanders et al. 2015) and Developmental Delay (DD) (Deciphering Developmental Disorders Study 2017). Similar to previous studies (An et al. 2018), genes were included that interact transcriptionally or physically with genes in these sets such as loci bound by the ASD gene product CHD8 (Cotney et al. 2015), and proteins that interact with Fragile X mental retardation protein FMRP (Darnell et al. 2011). We quantified OCRs nearby a gene using the inverse distance from the TSS to the closest OCR and used the shortest of these distances if the gene has multiple TSSs. We then used OCR accessibility annotations in cortex versus basal ganglia (or both) to test if each neurodevelopmental disorder gene set was more associated with each brain region than expected by chance. Liver (Subramanian et al. 2005) and olfactory receptor (Rouillard et al. 2016) gene sets were used as negative controls. We also compared psychiatric disorder genes to a size-matched random set of genes expressed across the brain. To test for region specificity, we also included sets of genes highly expressed in cortex but not in basal ganglia, and vice versa.

FMRP targets and DD genes were significantly enriched in both cortex and basal ganglia, while ASD genes were significantly enriched in cortex. CHD8 targets and negative controls showed no enrichment. Highly expressed genes in cortex were enriched near cortex OCRs, while highly expressed genes in basal ganglia were enriched near both cortex and basal ganglia OCRs, though with a much larger odds ratio in the latter. We repeated this analysis for specific cortical and basal ganglia sub-regions and found neurodevelopmental disorder gene sets enriched in specific sub-regions (Figure S6B,C). Table S6B lists all region-specific OCRs proximal to neurodevelopmental disorder genes.

### Function altering *de novo* point mutations in pREs

To further investigate the potential roles of pREs in human disorders, we studied *de novo* variants identified by whole-genome sequencing of 1,902 quartet families (An et al. 2018), composed of an individual diagnosed with ASD, an unaffected sibling and both parents. The category-wide association study (CWAS) framework (Werling et al. 2018) divides the genome into over 50,000 categories defined by functional annotations, conservation across species, gene-defined regions, or proximity to genes implicated in ASD, each of which is tested for enrichment of *de novo* variants in cases vs. controls. Correcting for the categories tested, no single noncoding category has reached significance. However, a *de novo* risk score (DNRS) to assess risk across multiple categories implicated promoter regions (An et al. 2018). Adding the pREs to the CWAS analysis did not yield a statistically significant result after multiple-testing correction. However, intronic pREs near ASD-associated genes showed a strong trend towards enrichment (Figure S7).

The gene *SLC6A1*, encoding the GABA transporter GAT-1, is associated with ASD (Sanders et al. 2015) and myoclonic atonic epilepsy/absence seizures with developmental delay (Heyne et al. 2018). *De novo* variants from two individuals with ASD but no seizures were identified by WGS (An et al. 2018) and mapped to an intronic pRE near *SLC6A1*. This pRE has increased chromatin accessibility in the basal ganglia compared to cortex (Figure 7A); the mouse homologue shows open chromatin in neonatal cortical layer 6, but not layer 5, neurons (Figure 7B). We tested the function of the two *de novo* variants in this pRE, and found they significantly reduced enhancer activity using a luciferase transcription assay in neuroblastoma cells (Figure 7D). We also tested a *de novo* point mutation in an intronic pRE of histone lysine-9 demethylase encoding gene *KDM4B* with cortex-specific chromatin accessibility (Figure 7C); a *de novo* ASD point mutation in this pRE also reduced luciferase transcription (Figure 7D). These applications illustrate the utility of pREs for identifying functional enhancers and enhancer mutations.

**Figure 7:**
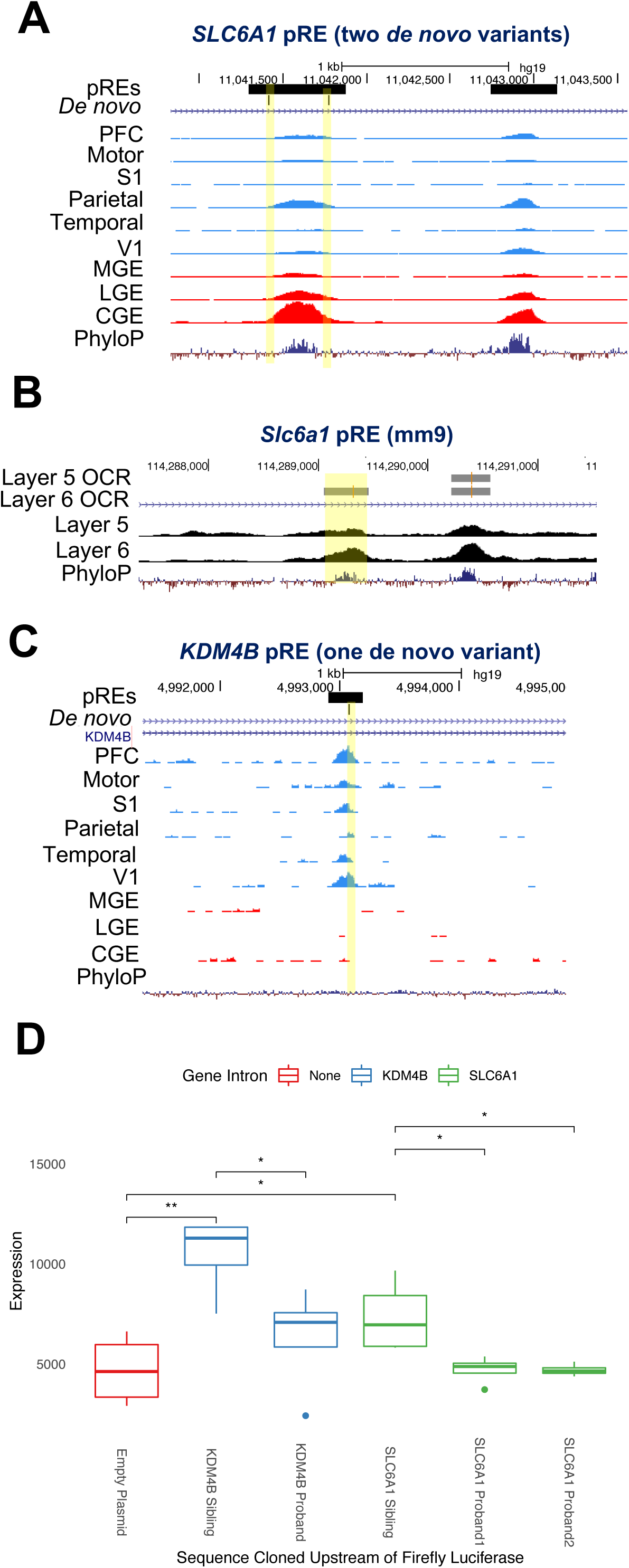
Functional *de novo* variants in pREs. 7A) ATAC-seq reads, combined across multiple samples per brain region, at ASD and epilepsy gene *SLC6A1*. The *SLC6A1* intronic pRE contains two *de novo* variants, highlighted in yellow, from separate probands. The *SLC6A1* pRE is called an OCR in all brain regions assayed except motor cortex and S1. Y-axis scale is 0 to 50. 7B) ATAC-seq reads at mouse gene locus *Slc6a1*, from purified populations of layer 5 and layer 6 neurons. Layer 5 and Layer 6 called OCRs are indicated. The homologous region to the human pRE containing two *de novo* mutations is highlighted yellow. 7C) The *KDM4B* intronic pRE contains a *de novo* variant from an ASD proband highlighted in yellow, and is an OCR in PFC, S1, and V1. Y-axis scale is 0 to 50. 7D) Mean firefly luciferase levels in human neuroblastoma cells, normalized to Renilla, testing enhancer activity of *SLC6A1* and *KDM4B* pREs and the functional effects of proband point mutations in the pREs on luciferase expression levels. The two pREs were cloned upstream of a minimal promoter and firefly luciferase ORF in the pGL4.23 vector and luciferase levels in whole cell lysates were measured four days after transfection. Error bars indicate standard error across four replicate experiments.

## Discussion

We generated an atlas of the chromatin accessibility landscape across nine regions of the mid-gestation human telencephalon and predicted a subset of open chromatin regions (18%) most likely to function as regulatory elements (pREs). A substantial proportion of these pREs show regional, temporal, and laminar differences in chromatin accessibility. Across these three axes, we linked pREs to differentially expressed genes by integrating single-cell RNA-seq data, and identified TFs that may bind pREs. We show that this data can be used to identify enhancers of genes associated with neurodevelopmental disorders, including a validated enhancer of *BCL11A*, and two *de novo* mutations in the same pRE in individuals with ASD that alter expression of *SLC6A1*. Regional differences in the activity of pREs containing ASD-associated variants provide insights into anatomical locations where the genes’ function may be critical during brain development.

### An atlas of candidate enhancers in the developing telencephalon

This annotated collection of ATAC-seq data from nine telencephalic regions and two cortical laminae has been integrated with single-cell RNA-seq analyses (Nowakowski et al. 2017) to identify elements that may drive transcriptional differences between developmental stages, cell types, brain regions, and cortical layers. The mid-fetal telencephalon undergoes numerous developmental processes including proliferation, neurogenesis, cell type specification, migration, integration into neural structures (cortex, striatum and pallidum), and the beginning of neuronal maturation (formation of dendrites, axon pathways, synapses); our gene ontology analyses suggest that individual pREs may regulate these processes. New technologies enabling simultaneous investigation of gene expression and chromatin accessibility (Cao et al. 2018) will allow direct interrogation of the chromatin dynamics that accompany neurodevelopmental processes at the single-cell level. Our data will serve to anchor forthcoming single-cell epigenomic analyses.

Transgenic mouse experiments have illustrated the exquisite spatiotemporal and cell type-specific activity of neurodevelopmental enhancers afforded by combinatorial binding of TFs expressed in graded, overlapping patterns in the brain (Silberberg et al. 2016; Pattabiraman et al. 2014; Visel et al. 2013; Erwin et al. 2014). TFs predicted to bind human pREs that specify pallial and subpallial structures include T-box family and Nkx family TFs, respectively, which have been shown in mice to specify cortical excitatory neuron and MGE-derived interneuron subtypes by binding at distal REs. Interestingly, the OCT4/SOX2 motif was enriched in basal ganglia specific pREs (Figure 3D), and the function of this composite motif has been demonstrated in a mouse MGE enhancer of *Tcf12* (Sandberg et al. 2018). Within pREs that were differentially accessible over cortical ages or cortical lamina, we also identified motifs for TFs that have been well-studied in mice, indicating their relevance to human brain development. The pREs provided here (Table S2) may be integrated with mouse ChIP-seq studies of these and other TFs to indicate which loci of interest may be functional enhancers in different regions of the developing human brain.

This pRE atlas can also be integrated with ATAC-seq data from other species. For instance, we integrated human ATAC-seq data with ATAC-seq data generated from layer 5 and 6 purified neurons of genetically modified mice. Using this approach, we attributed chromatin accessibility in human deep layer pREs to specific cortical neuronal subtypes (Figure 5). The pREs provided here are also a valuable resource for curating a subset of loci accessible in mouse cell types which are likely to be conserved neurodevelopmental enhancers in human.

The database presented here is accessible and searchable by area of interest, and contains the following fields: (1) *Brain Region*. Table S2A organizes all pREs by the brain region in which they overlap OCRs, and notes the pREs specific to basal ganglia, cortex, and pREs specific to just one telencephalic subregion. Table S3 lists pREs proximal to genes differentially expressed between MGE, PFC, and V1, whose chromatin accessibility is specific to each pairwise comparison; (2) *Cortical Laminae*. pREs for upper and deep layer cortical neurons in fetal PFC, which are enriched for upper and deep layer TF motifs, are listed in Table S5. This also notes which pREs are conserved OCRs to mouse layer 5 and 6 cortical neurons; (3) *Cortical developmental stage*. pREs for early and late mid-fetal frontal cortex are listed in Table S4; (4) *Cell Type.* Table S6C provides genes that are expressed exclusively in particular cell types in the developing telencephalon and the region specific OCRs that are found near those loci; (5) *Gene and Locus.* Table S2A lists the upstream and downstream genes of all predicted pREs, (6) *Human disorders.* Table S6B provides region specific OCRs that are proximal to ASD-associated genes, developmental delay genes, FMRP targets, and CDH8 targets. (7) *De Novo Noncoding Variant.* Table S7 lists pREs that contain more than one ASD patient mutation (An et al. 2018).

We emphasize that pREs are predictions based on enhancers validated via transgenic mouse assays and described by numerous diverse sequence and functional genomics features. We validated enhancer activity of twelve pREs by luciferase and transgenic assays (Figures 2, 6, and 7). However, further work is needed to test their function *in vivo* and investigate whether the activity of pREs can be therapeutically modulated, as has been done for obesity using CRISPRa (Matharu et al. 2019) to compensate for haploinsufficiency or over-expression. In the future, our machine learning approach could be applied to collections of in vitro validated enhancers.

### Implications for human genetics and disorders

This atlas has various implications to the field of neurogenetics. One challenge for the field is the limited functional annotation of the noncoding genome where many risk alleles are thought to be located. We address this by providing functional annotation of OCRs and pREs in the noncoding genome of the developing human brain. Another challenge is predicting the impact of noncoding genetic variation on brain development. This atlas provides a roadmap for identifying the telencephalic regions, cortical neuron cell types, time periods, and TF networks of the developing mid-gestation telencephalon that may be impacted by genetic variation in human traits and disorders.

Identifying the regulatory elements of genes implicated in neurodevelopmental disorders is essential for elucidating the transcriptional circuitry organizing their expression. Intronic pREs of ASD genes that contain case mutations, such as the intronic *SLC6A1* pRE (Figure 7), may be important developmental enhancers for those genes and may provide a mechanism to modify expression as a future therapy (Matharu et al. 2019). Defining when and where these pREs are active may point to which brain regions and cell types underlie neurodevelopmental disorders.

Similarly, identifying the downstream REs controlled by ASD risk genes, many of which encode transcription regulators or chromatin modifiers (De Rubeis et al. 2014; Cotney et al. 2015), is critical in understanding how mutations alter expression programs impacting brain development and function. Indeed, ASD genes and FMRP targets appear to be under precise epigenetic control, as they are significantly enriched for region-specific OCRs (Figure 6C, Figure S6B,C). These OCRs may be sensitive to epigenetic alterations that occur in the brain upon loss of function of ASD genes (Katayama et al. 2016; Cotney et al. 2015) or FMRP (Korb et al. 2017), and may indicate which brain regions are specifically impacted by the resulting chromatin changes.

The chromatin accessibility data curated here will be a useful resource for studies elucidating the transcriptional networks dysregulated in human disorders. Our study has other clinical implications beyond the importance for interpreting the significance of genetic variation in candidate REs. For example, a recent postmortem analysis of ASD discovered that alterations in upper layer neuron gene expression are correlated with clinical severity (Velmeshev et al. 2019); thus, upper cortical layer pREs identified here may be important targets for therapy.

## Acknowledgments

This work was supported by the research grants to JLR from: Nina Ireland, NINDS R01 NS34661 and to KSP from: NIMH R01 MH109907 and NIMH U01 MH116438. ATAC-seq and ChIP-seq datasets for early neural differentiation timepoints were provided by the Ahituv lab (Inoue et al. 2018).

## Author Contributions

Conceptualization, EMP, SW, KSP, JLR; Investigation, EMP, SW; Methodology, EMP, SW, PP, RT, KSP; Validation, EMP, FB; Software, SW, PP, RT; Formal Analysis, EMP, SW, PP, RT; Resources, TN, SS, JLR, KSP; Data Curation, SW, SS; Writing – Original Draft Preparation, EMP, SW, PP, KSP, JLR.; Writing – Review & Editing, EMP, SW, PP, SS, MWS, KSP, JLR; Visualization, EMP, SW, PP; Supervision, SS, MWS, KSP, JLR; Project Administration, EMP, SW, KSP, JLR.; Funding Acquisition, MWS, KSP, JLR.

## Supplemental Figure Legends

**Figure S2:**
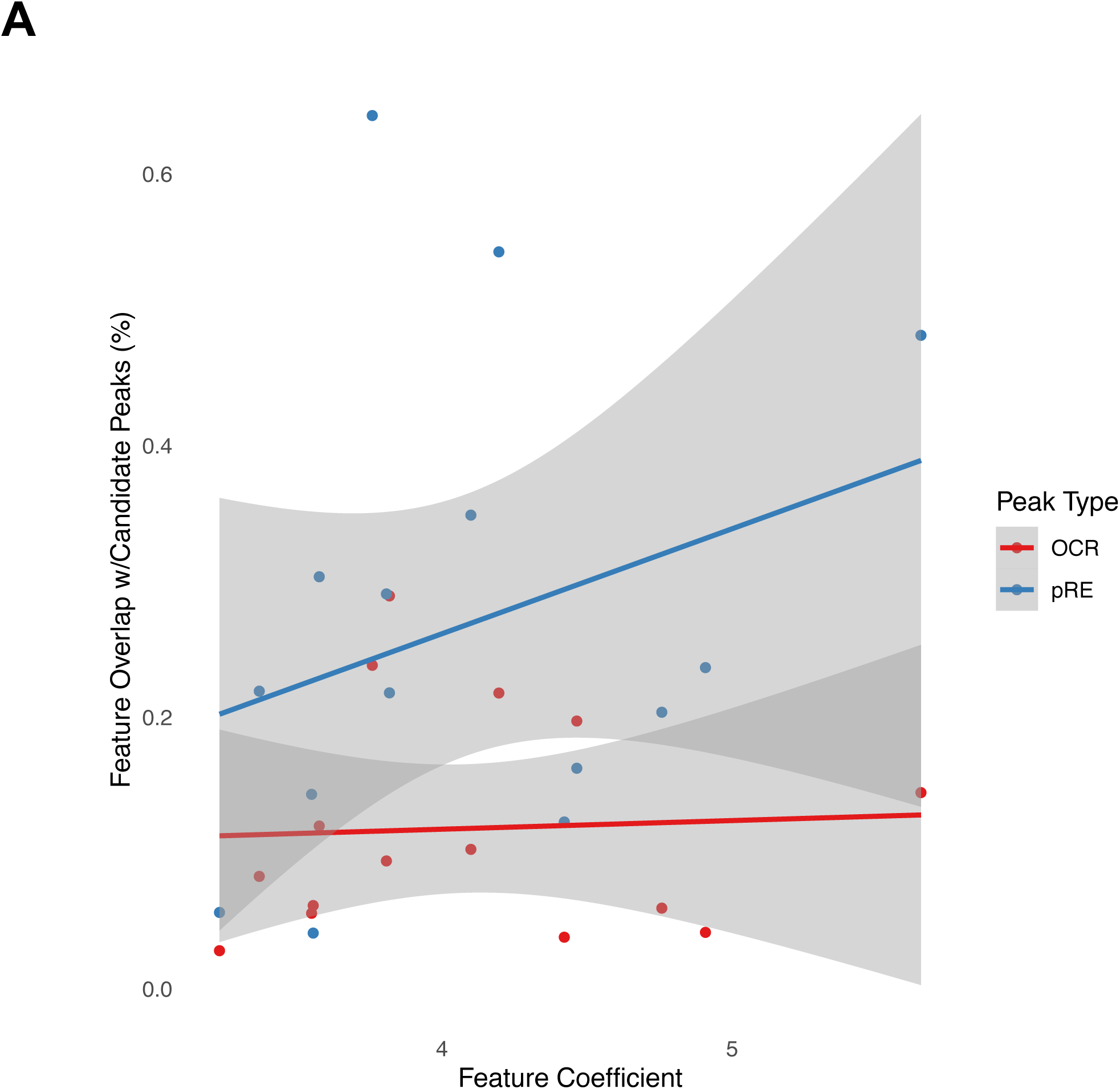
Predicting neurodevelopmental enhancers. Features with the largest 15 coefficients selected by a penalized linear model for predicting neurodevelopmental enhancers, plotted against the percentage of OCRs and pREs overlapping peaks from a given dataset. pREs in general have greater overlap with the model’s most important features and, unlike OCRs, have a modest correlation between coefficient and peak overlap. This provides additional intuition for why the model selected certain features.

**Figure S3:**
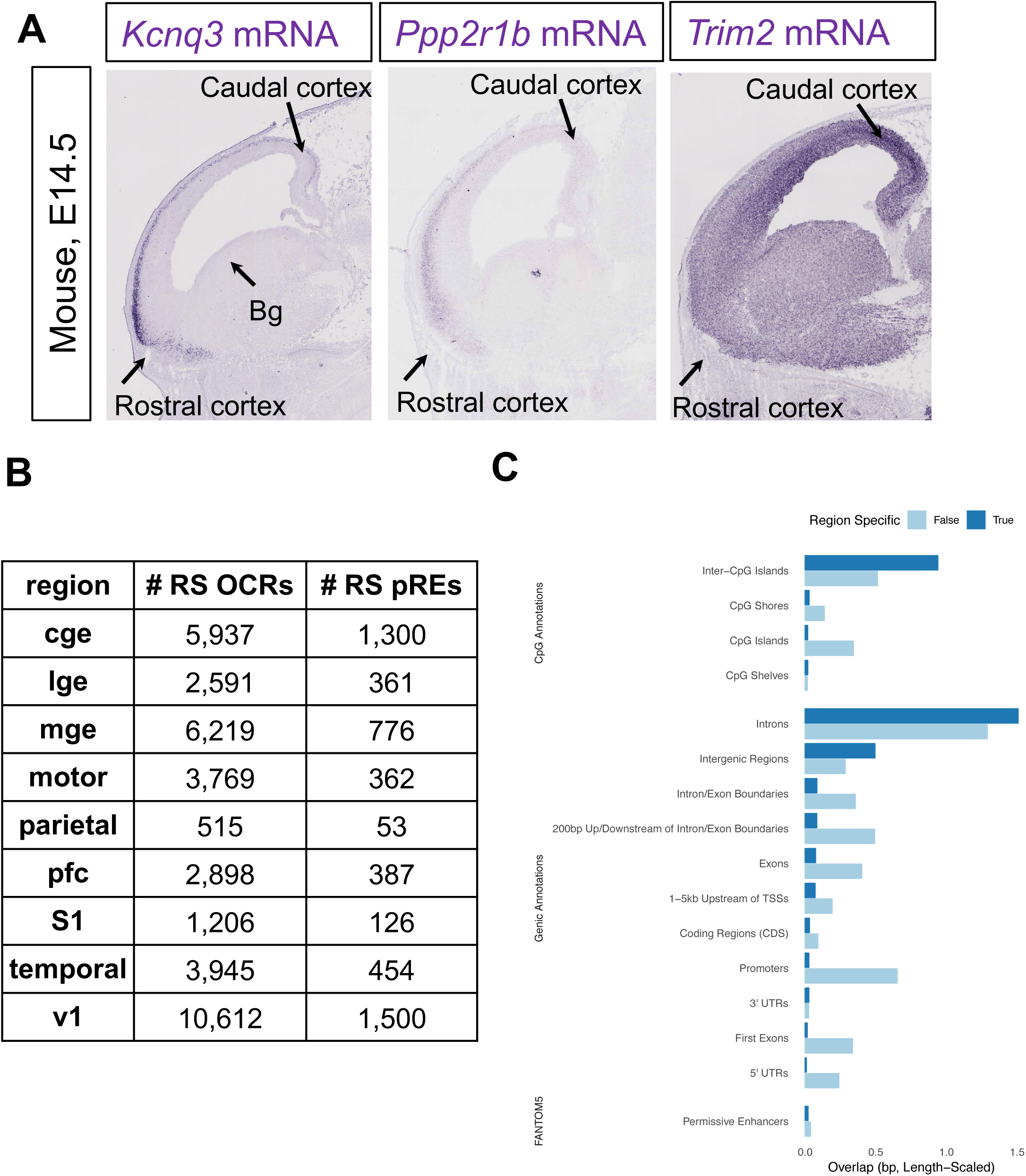
Regional differences in chromatin accessibility at pREs. S3A) *In situ* hybridization for *Kcnq3, Pppp2r1b*, and *Trim2* mRNA (purple) in wildtype murine embryos at E14.5, from Eurexpress database (www.eurexpress.org) (Diez-Roux et al. 2011). *Kcnq3* is expressed in the cortex (both rostral and caudal) and not the basal ganglia (bg), *Pppp2r1b* is expressed in rostral cortex, and *Trim2* is expressed in caudal cortex. S3B) Summary of region-specific OCRs and pREs identified in this study. S3C) Annotation of genomic features at region-specific pREs compared to those shared across multiple brain regions.

**Figure S4:**
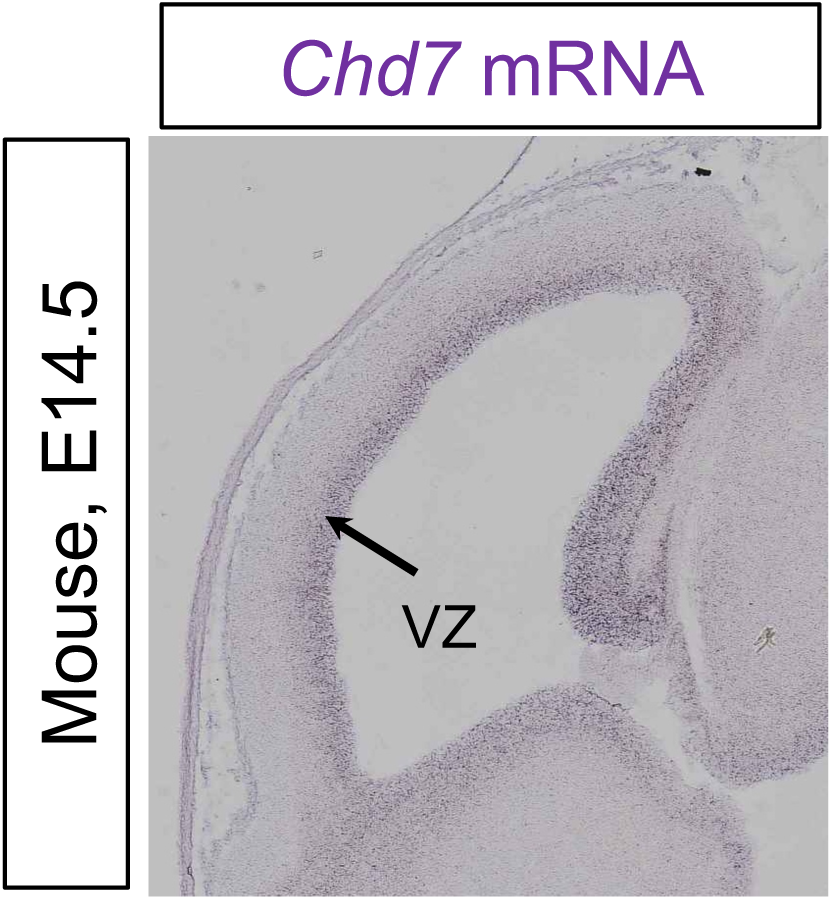
*In* s*i*tu hybridization for *Chd7* mRNA (purple) in wildtype murine embryos at E14.5, from Eurexpress database (www.eurexpress.org) (Diez-Roux et al. 2011). Expression in the ventricular zone (VZ) of cortex is indicated.

**Figure S5:**
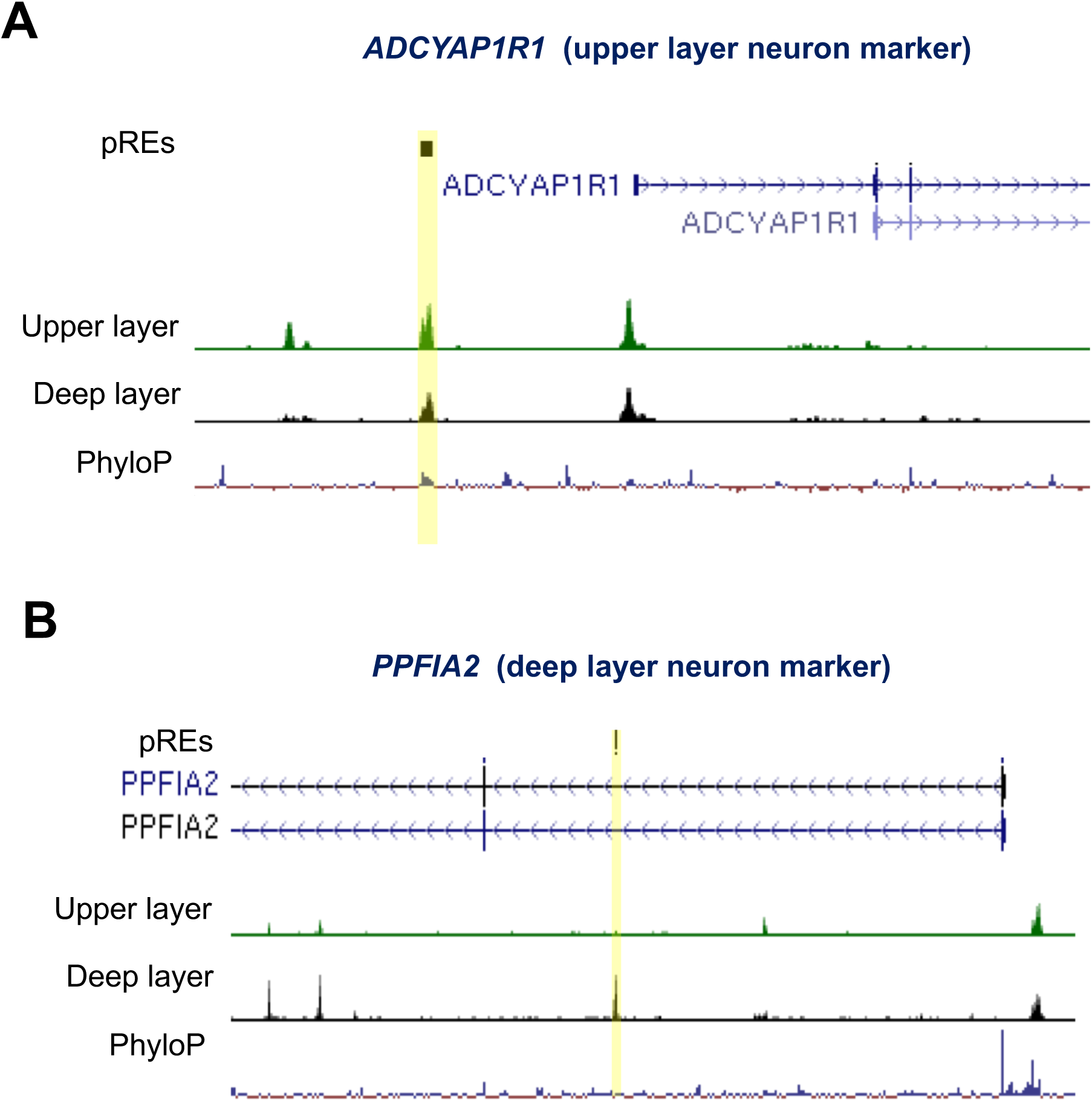
ATAC-seq reads combined from multiple samples from upper layer and deep layer microdissections of PFC at 18gw and 19gw, mapped to A) upper and B) deep layer marker genes. Layer-specific pREs are indicated in yellow. Y-axis scale is 0 to 50.

**Figure S6:**
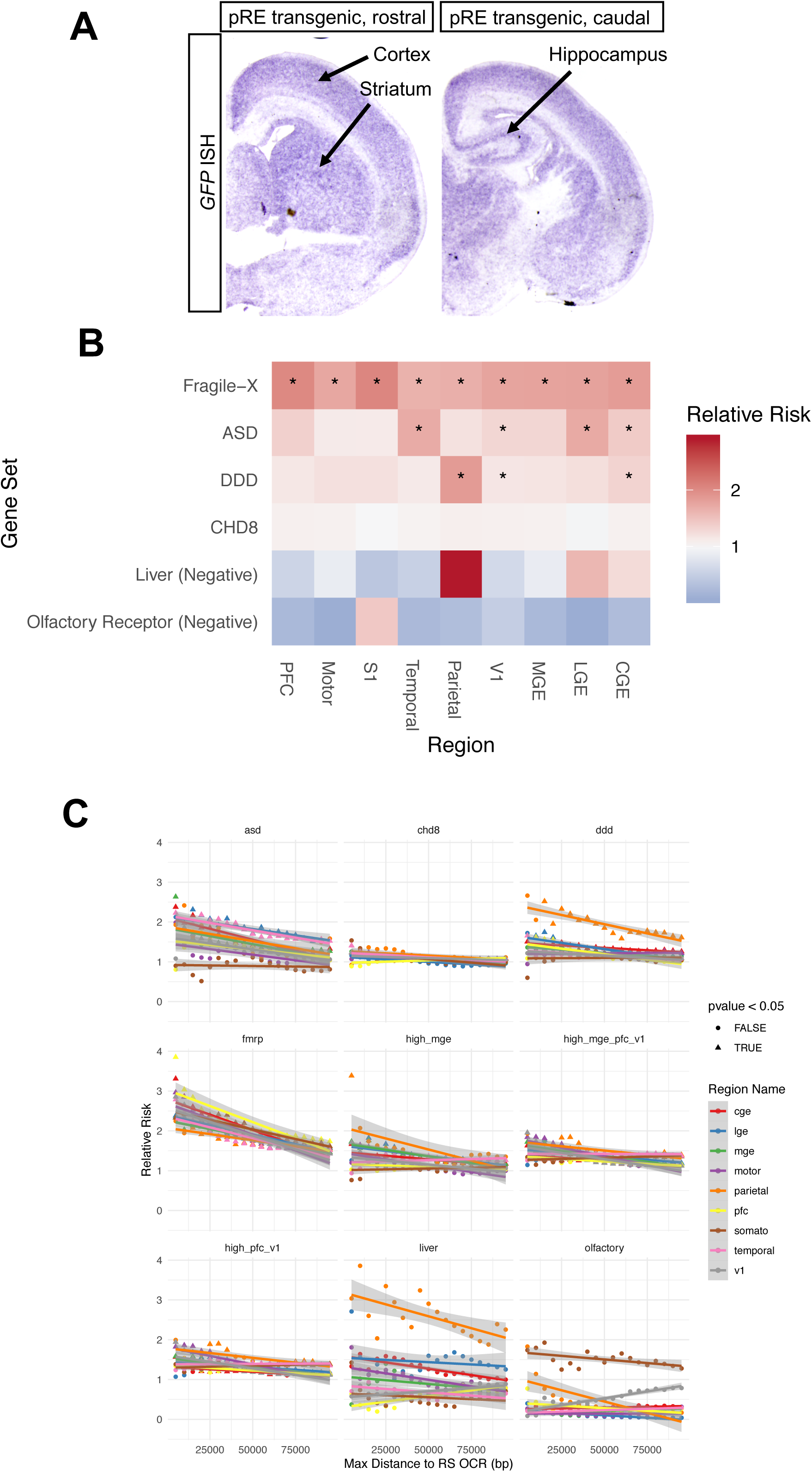
S6A) Postnatal day 2 sections of *BCL11A* enhancer transgenic mouse, a separate founder line than the one shown in Figure 6. GFP reporter expression in purple (RNA in situ hybridization for GFP). Cortical, striatal, and hippocampal expression is indicated. S6B) Enrichment analysis of genes with a TSS proximal (within 50 kb) to region-specific OCRs. ASD genes (Sanders et al. 2015), biological targets of Fragile X Mental Retardation protein (FMRP) (Darnell et al. 2011), biological targets of ASD gene CHD8 (Cotney et al. 2015), and developmental delay (DDD) genes (Deciphering Developmental Disorders Study 2015) were compared to liver (Subramanian et al. 2005) and olfactory receptor (Rouillard et al. 2016) gene sets. S6C) Enrichment analysis of genes having a TSS within a set maximum distance of region-specific OCRs, using multiple distance thresholds from 5 kb to 100 kb in increments of 5kb. A general trend for higher enrichment is seen for more proximal distance thresholds, though these points are not always statistically significant (indicated by point shape) due to the reduced number of RS OCRs for testing.

**Figure S7:**
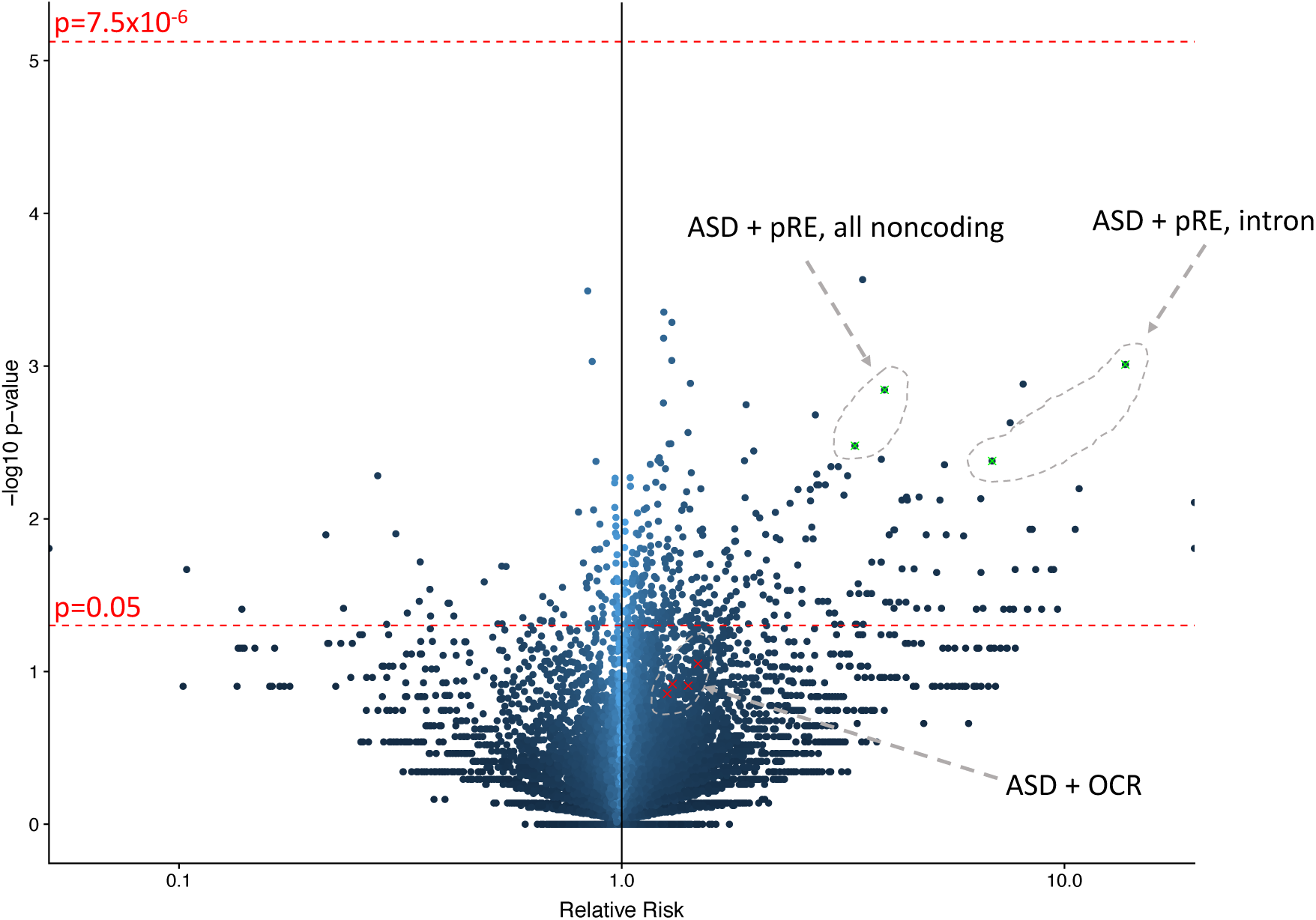
Category wide association study (CWAS) on *de novo* ASD mutations from 1,902 quartet families (An et al. 2018) using genome annotations such as conservation scores, gene proximity to neurodevelopmental disorder genes, OCRs and pREs. Although pRE-containing categories did not cross the threshold for statistical significance (p-value = 7.5*10^-6^ after Bonferroni correction for 6,711 effective tests), they had higher relative risks (labeled and highlighted in grey; ASD=ASD gene locus) than the same categories using OCRs rather than pREs. This demonstrates the value of our machine-learning approach for honing in on a subset of OCRs related to disease.

## STAR Methods

The hg19 reference genome was used in all analyses due to the integration of many diverse datasets, with GRCh38 coordinates lifted over to hg19 where necessary. Analyses were conducted using python, pandas (McKinney 2012), R (R Core Team 2018), bioconductor (Huber et al. 2015), and bedtools2 (Quinlan and Hall 2010).

### ATAC-seq library generation from human samples

De-identified tissue samples were obtained with patient consent in strict observance of the legal and institutional ethical regulations. Protocols were approved by the Human Gamete, Embryo, and Stem Cell Research Committee, the institutional review board at the University of California, San Francisco.

Fresh fetal brain samples were obtained in accordance with etc etc and were transported in freshly made Cerebral Spinal Fluid on ice (CSF). All dissections and ATAC-seq experiments were performed within two hours of tissue acquisition. Dissections of each brain region included the entire telencephalic wall, from the ventricular zone to the meninges, except for the upper and deep layers where the cortical plate from the PFC was microdissected under a microscope. From each dissection, intact nuclei were isolated by manually douncing the tissue twenty times in 1mL Buffer 1 (300mM sucrose, 60mM KCl, 15mM NaCl, 15mM Tris-HCl, pH 7.5, 5mM MgCl2, 0.1mM EGTA, 1mM DTT, 1.1mM PMSF, Protease inhibitors) on ice using a loose pestle douncer, and then lysed on ice for 10 minutes after adding 1mL Buffer 2 (300mM sucrose, 60mM KCl, 15mM NaCl, 15mM Tris-HCl, pH 7.5, 5mM MgCl2, 0.1mM EGTA, 0.1% NP-40, 1mM DTT, 1.1mM PMSF, Protease inhibitors). During this ten minutes, nuclei were counted using trypan blue and 50,000 nuclei were spun down at 7,000rpm for ten minutes at 4C. Nuclei were resuspended in 25uL Tagmentation buffer, 22.5 uL Nuclease Free H20, and 2.5 uL Tagmentation Enzyme from Illumina DNA Kit (number), gently mixed, and placed in 37C water bath for thirty minutes. THe tagmentation reaction was stopped by MinElute PCR purification and DNA was eluted in 10uL Nuclease Free water. ATAC-seq library generation was performed using Illumina barcode oligos as described (Buenrostro et al 2015), for 8-11 cycles PCR. The number of cycles was empirically determined for each library. Libraries were bioanalyzed using Agilent High Sensitivity DNA Kit, pooled together and sequenced on Hiseq 2500 using paired end 50 bp sequencing.

### ATAC-seq library generation from mouse samples

Postnatal day 2 mice from Ntsr1-CRE or Rbp4-CRE (Gong et al. 2003) male mice crossed to Intact fl/fl female mice were sacrificed and the PFC was dissected. Nuclei from individual mouse brains were isolated as above using 1mL Buffer 1 (see above), douncing gently with loose pestle on ice, and lysing in Buffer 2. After centrifugation, nuclei were resuspended in PBS with FCS and taken to FACAria fluorescent cell sorter. 50,000 GFP positive nuclei were isolated, spun down, and resuspended in Tagmentation reaction, and placed in 37C water bath for thirty minutes. Reaction was stopped by MinElute PCR purification and DNA was eluted in 10uL Nuclease Free water. ATAC-seq libraries were prepared, bioanalyzed on Agilent High Sensitivity DNA kit, and sequenced on Hiseq 2500 using paired end 100bp sequencing.

### Peak Calling

Paired-end reads were aligned and peaks were called using the ENCODE ATAC-seq pipeline with default parameters (Lee et al. 2016). The pipeline produces multiple sets of peak calls, including those generated by macs2 (Zhang et al. 2008) and the Irreproducible Discovery Rate (IDR) package for R (Q. Li 2014). The latter was used to select a smaller, more confident set of peaks that are likely to be consistent across biological replicates (Q. Li et al. 2011). Peaks overlapping ENCODE blacklisted regions were removed (Amemiya, Kundaje, and Boyle 2019). The pipeline created pseudo-replicates where a portion of reads are held-out in order to estimate a threshold for calling consistent peaks.

### Peak Merging

For each (region, timepoint) combination, peaks separated by up to 100 bp were merged into a single peak to reduce variation in peak calls attributable to coverage differences (bedtools merge -d 100). The union of peaks across all samples was intersected with itself to identify peaks overlapping by at least 75% (bedtools intersect -f 0.75 -e). These overlaps were converted to a matrix with peak coordinates as rows and (region, timepoint) combinations as columns. A (region, timepoint) column was assigned a 1 when one of its peaks overlapped a row’s peak, otherwise a 0 was assigned.

To identify region-specific peaks, columns corresponding to the same brain region were merged: a 1 was assigned if a peak was called for any timepoint in that region, otherwise a 0 was assigned. This resulted in a new matrix with peaks as rows and regions as columns. Region-specific peaks were then those peaks with a 1 in a single column (region) and 0s in all other columns (regions).

### Disease Gene Enrichment in Marker Genes, RS OCRs

Disease gene sets (An et al. 2018) and single cell marker genes (Nowakowski et al. 2017) were downloaded. Marker genes were manually associated with the appropriate brain regions of our samples. Genes were annotated with transcription start site(s) from GENCODE v19 (Frankish et al. 2019). Logistic regression (R’s glm function with family = binomial) was performed on each combination of gene set and region using the inverse distance from a gene to the closest regional OCR as a covariate, and the presence or absence of the gene in the set as the dependent variable. Distance was capped at 1 megabase. One-sided p-values were computed from glm’s t-statistic using R’s pnorm function with lower_tail = False. Negative control gene sets were also used including liver (Subramanian et al. 2005), olfactory receptor (Rouillard et al. 2016), and a size-matched set of genes expressed in whole brain (Hawrylycz et al. 2012). To increase power for disease gene set testing, peaks were combined into cortex- or basal ganglia-specific OCRs, while single cell marker gene sets were tested in sample-matched regions.

### Regulatory Element Prediction

Elastic Net and Random Forest classifiers (Pedregosa et al. 2011) were trained on and generated predictions for OCRs merged across all brain regions. Training labels were generated using OCRs in combination with VISTA enhancers. OCRs were labeled positive if they overlapped validated developmental brain enhancers, or labeled negative if they overlapped validated non-brain enhancers or enhancers that failed to validate. Features were generated for each OCR including 5-mers (counts of all 5 consecutive base pairs occurring within the peak), binary overlaps with ChIP-seq peaks and average peak methylation from Roadmap Epigenomics (Roadmap Epigenomics Consortium et al. 2015), binary overlaps with either fragment of statistically significant Hi-C loops (Won et al. 2016; Rao et al. 2014), and average evolutionary conservation across the region (Siepel et al. 2005; Pollard et al. 2010). For each peak without a training label, a continuous-valued prediction was generated corresponding to the model’s confidence that the OCR is a developmental brain enhancer. OCRs were considered pREs if either algorithm’s prediction was above 0.5, representing 19,151 out of 103,829 OCRs (18.4%) after merging across samples and excluding promoter overlaps (1500 bp upstream, 500 bp downstream from a GENCODE v19 TSS (Frankish et al. 2019)).

### Luciferase Assay

The following primers were used to amplify pREs from human genomic DNA, then cloned into the minimal promoter pGl4.23 luciferase vector (Promega) using SacI and XhoI restriction sites (underlined) in the vector’s multiple cloning site.

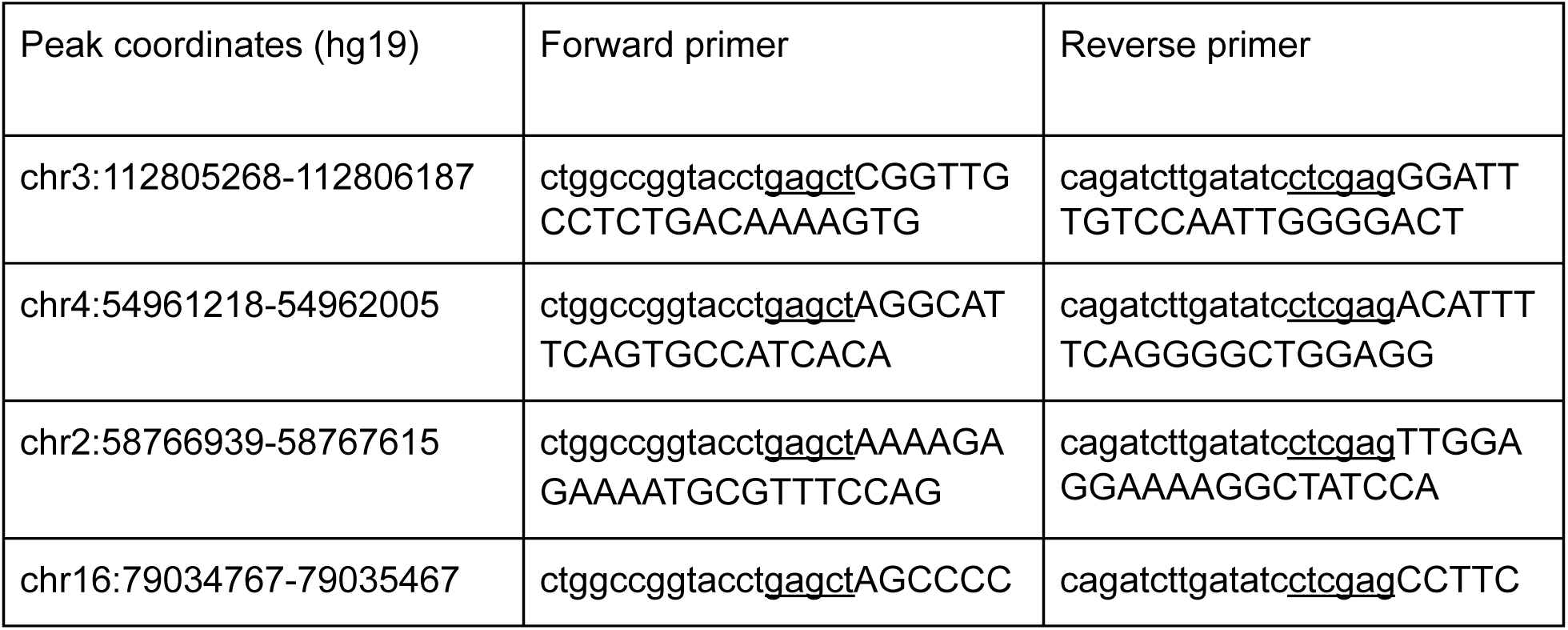
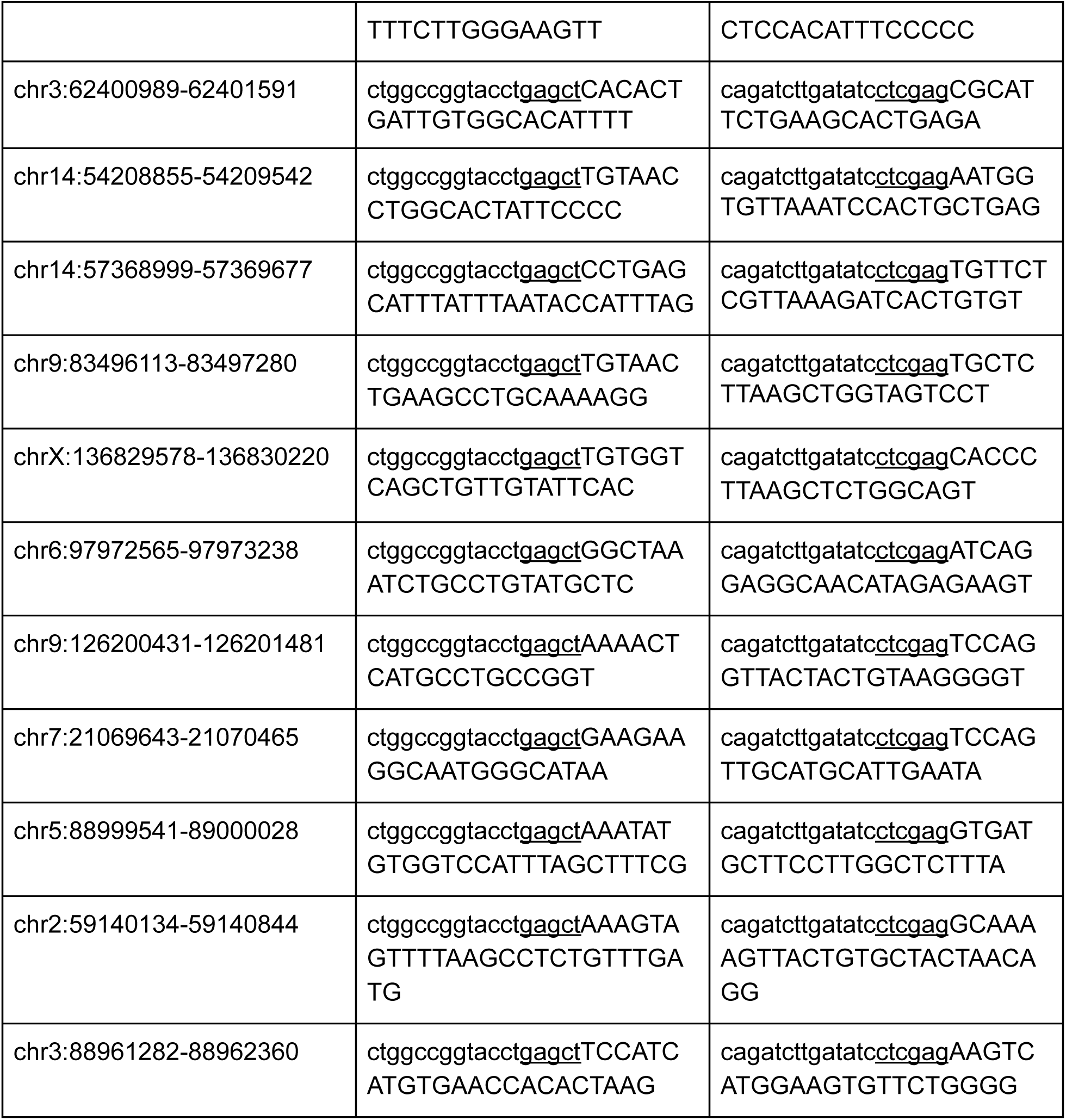

Human brain-derived neuroblastoma cells BE(2)-C from ATCC were passaged three times and grown to confluency in a 1:1 mixture of Eagle’s Minimum Essential Medium and F12 Medium with 10% fetal bovine serum. Confluent cells were transfected in four 96-well plates with luciferase vectors (predicted enhancer-pGL4.23 or empty vector pGL4.23) and renilla vector. Two days later, cells were lysed and luciferase levels detected using the Promega dual reporter luciferase assay kit. Luciferase levels were normalized to Renilla and averaged across four replicate experiments.

### Transgenic enhancer mouse generation and analysis

To clone the putative BCL11A enhancer into the CT2IG vector, the following primers were used:

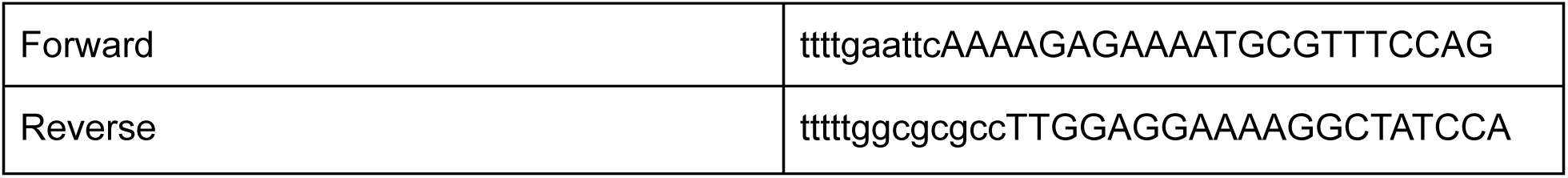

A 960 bp amplicon from human genomic DNA was gel purified, digested with EcoRI and AscI restriction enzymes, and ligated into linearized CT2IG vector upstream of a minimal hsp68 promoter, tamoxifen inducible Cre, IRES, and green fluorescent protein (GFP). Sanger sequencing confirmed insertion of the pRE into the CT2IG vector and linearized vector was submitted for pronuclear injection of CD1 mice at the Gladstone Transgenic Core Facility. Founders were screened for the transgene using the following genotyping primers:

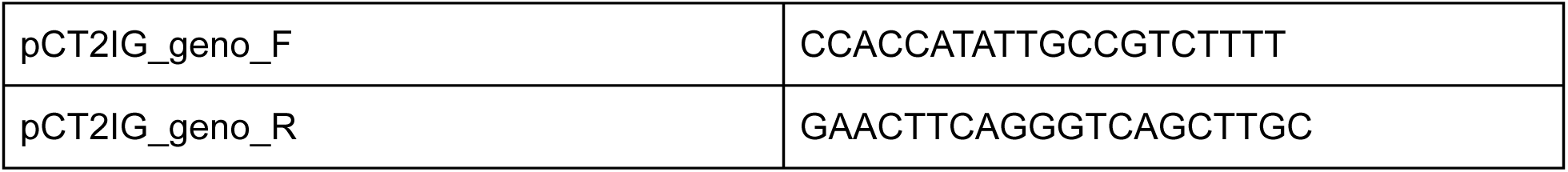

Three positive founders were bred to CD1 wildtype mice, and F1 generation mice were analyzed at postnatal day 2. Postnatal day 2 mice were perfused with 4% paraformaldehyde, whole brain was dissected, and postfixed overnight in 4% paraformaldehyde, and transferred to 30% sucrose overnight. 20 micron thick cryosections were obtained and in situ hybridization using GFP RNA probe was performed as described (Sandberg et al. 2018). In situs were developed at 37C and were imaged two days later.

### Analysis of Relative Risk of *De Novo* ASD Mutations in Enhancers

Annotated *de novo* ASD mutations (An et al. 2018) were intersected with OCRs and pREs to annotate each mutation as having those as functional annotations. CWAS was conducted as described in An et al., using OCRs and PREs in addition to their noncoding annotation categories.

### Testing function of *de novo* variants in pREs

The following primers were used to amplify pREs in ASD gene introns from human genomic DNA. They were then cloned into the minimal promoter pGl4.23 luciferase vector (Promega) using SacI and XhoI restriction sites (underlined) in the vector’s multiple cloning site.

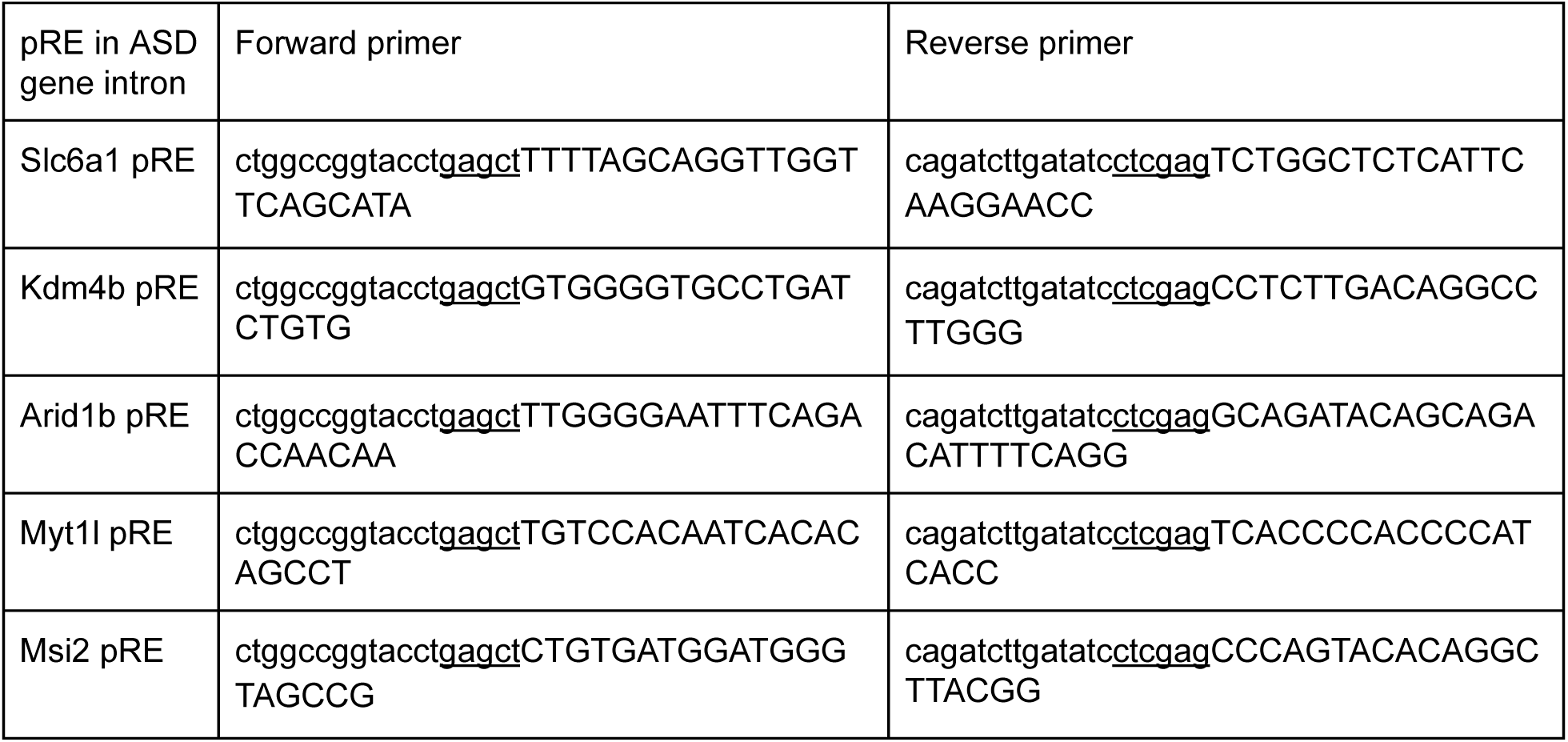

The above pREs were tested for enhancer activity by luciferase assay in BE(2)-C as described above. To introduce point mutations (Table S7) into these pREs, we used site directed mutagenesis. Phusion PCR was performed using the Slc6a1-pGl4.23 and Kdm4b-pGl4.23 vectors as template and the following two primers to introduce the *de novo* point mutation (capitalized) into the pRE sequence. All variants were taken from An et al. supplemental data.

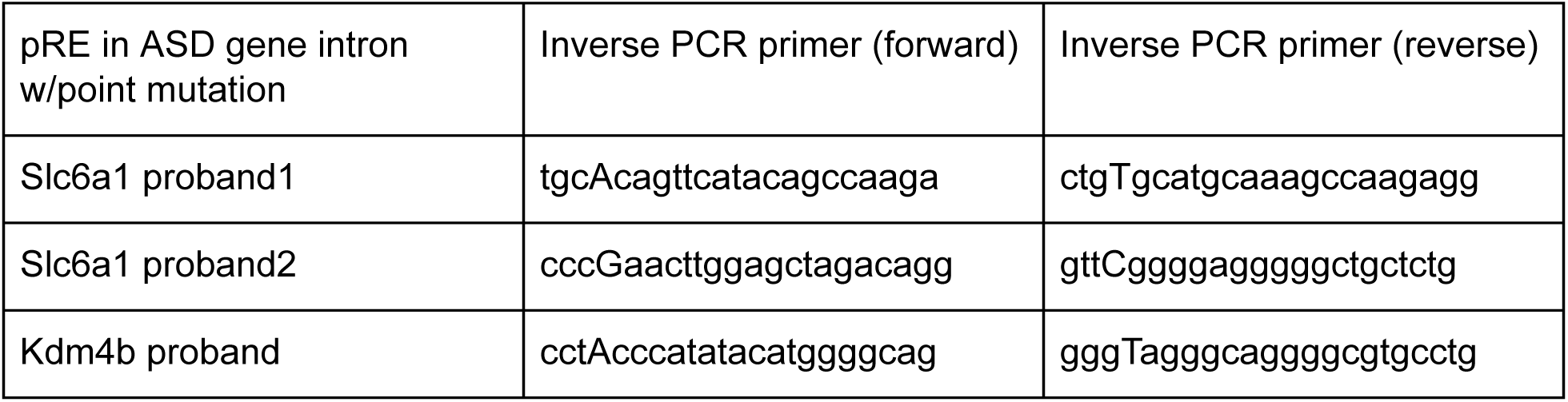

Luciferase levels were compared between empty vector, unmutated (sibling) pRE, and pREs containing the above mutations. Four replicate experiments inBE (2)-C cells were performed and analyzed as described above.

### Motif Enrichment

The findMotifsGenome.pl script provided by Homer (Heinz et al. 2010) was used to identify TF motifs enriched in pREs that overlapped OCRs in one or more brain regions, depending on the analysis. The set of genomic regions used as foreground and background are provided below:

#### Cortex versus Basal Ganglia

Foreground: OCRs present in at least one of the cortical regions (pfc, motor, parietal, somato, temporal) and overlapping a pRE

Background: OCRs absent in all basal ganglia regions (cge, mge, lge) and overlapping a pRE

#### Cortex

Foreground: OCRs present only in the cortical region of interest (pfc, motor, parietal, somato, or temporal) and overlapping a pRE

Background: OCRs open in more than one cortical region and overlapping a pRE

#### Basal Ganglia

Same as for cortex, but with basal ganglia regions (cge, mge, lge)

#### PFC Deep Layer versus Upper Layer

Foreground: OCRs present only in PFC deep layer neurons and overlapping a pRE

Background: OCRs present only in PFC upper layer neurons and overlapping a pRE

#### Early versus Late Frontal Cortex

Foreground: OCRs present in at least one frontal cortex region (pfc, motor) at 14gw and overlapping a pRE

Background: OCRs absent in all frontal cortex regions at 19gw

### Association of Changes in Gene Expression with Chromatin State

Single cell RNA-seq data was downloaded for mge, pfc, and v1 brain regions (Nowakowski et al. 2017). MAST (Finak et al. 2015) identified differentially expressed genes between each pair of brain regions (V1 vs. PFC, V1 vs. MGE, PFC vs. MGE, FDR 5%). For each pair, differentially accessible pREs were associated with nearby genes using GREAT (McLean et al. 2010). Proximal associations used default parameters (5 kb upstream and 1 kb downstream of the TSS) while distal associations were restricted to a 100 kb window around the TSS. For a pair of brain regions, R’s fisher.test function estimated the odds ratio of differentially expressed genes over non-differentially expressed genes being linked to a uniquely open chromatin region. This same analysis was performed using genes differentially expressed between early and late frontal cortex timepoints, and again for genes differentially expressed between PFC upper and deep layers. These analyses were performed independently for OCRs and pREs.

### Peak Annotation

Genomic annotations were computed (Cavalcante and Sartor 2017) for both region-specific and non-specific pREs. The amount of each overlap (in bp) was scaled by the total length of the pRE.

